# Oxytocin neurons signal state-dependent transitions from rest to thermogenesis and behavioral arousal in social and non-social settings

**DOI:** 10.1101/2024.10.22.619715

**Authors:** Morgane Vandendoren, Jason G. Landen, Joseph F. Rogers, Samantha Killmer, Baizar Alamiri, Celeste Pohlman, Glenn J. Tattersall, Nicole L. Bedford, Adam C. Nelson

**Affiliations:** Department of Zoology and Physiology, University of Wyoming, Laramie, WY, USA; Department of Biomedical Sciences, Creighton University School of Medicine, Omaha, NE, USA; Department of Biological Sciences, Brock University, St Catharines, Ontario, Canada

**Author notes:** Corresponding author: Adam C Nelson. These authors contributed equally (Morgane Vandendoren, Jason G. Landen, Joseph F. Rogers).

## Abstract

Core body temperature (Tb) is defended within narrow limits through thermoregulatory behaviors like huddling, nesting, and physical activity as well as autonomic responses like brown fat thermogenesis. While Tb displays regulated fluctuations across different behavioral states and rest/arousal cycles, the neural control of these transitions is poorly understood. Here, we investigate the relationship between oxytocin neurons of the paraventricular hypothalamus (PVN^OT^) and behavioral and autonomic thermoeffector pathways across physiological states in mice. First, we show that PVN^OT^ neurons are activated during social thermoregulation. We then demonstrate that *in vivo* PVN^OT^ calcium dynamics align with transitions from rest to thermogenesis and behavioral arousal. Counter to our initial hypothesis, these dynamics were observed in both social and non-social contexts. Using a computer vision model to track thermoeffector pathways, we demonstrate that precisely timed stimulation of PVN^OT^ neurons during low-Tb resting states increases thermogenesis followed by behavioral arousal. We therefore suggest a model in which PVN^OT^ neurons facilitate homeostatic state-dependent transitions in thermo-behavioral states.

## Introduction

Maintaining a relatively constant core body temperature (Tb) is a vital homeostatic need. While excessive deviations (e.g. >2°C) from the typical mammalian Tb of approximately 37°C can be harmful^1^, Tb normally fluctuates within a narrow range (typically 0.5 - 1°C) across sleep-wake cycles and behavioral states ^2–4^. These fluctuations often exhibit bimodal or multimodal distributions, indicating distinct thermal states. Additionally, Tb variations are increasingly understood not as passive outcomes of animal physiology, but as regulated, brain-initiated transitions between defended Tb “balance-points” ^2,5–7^. For example, the transition from the rest to the active balance-point involves modulation of both autonomic and behavioral thermoeffector pathways (i.e., regulated heat loss and production) to meet the energetic demands of movement and arousal ^2,7,8^. However, the neural mechanisms that govern these physiological transitions remain incompletely understood ^8,9^.

Thermoregulation is orchestrated by the integration of autonomic and behavioral effectors. Autonomic outputs, such as brown adipose tissue (BAT) thermogenesis and vasomotor tone, are well-characterized at the circuit level. This pathway integrates sensory signals from peripheral thermosensors in the preoptic area of the hypothalamus (POA), then relays this information to the dorsomedial hypothalamus (DMH), and then to sympathetic efferent neurons of the rostral medullary raphe (rMR) ^5,10^. By contrast, the neuronal substrates of behavioral thermoregulation—such as huddling, nesting, or physical activity ^7,11–13^—and how they integrate with autonomic outputs are less well defined. Although recent studies have made progress in identifying brain regions underlying some of the associated neural pathways ^14–17^, much remains unknown about how animals behaviorally thermoregulate.

Behavioral thermoregulation strategies are naturally organized into alternating sequences across bouts of arousal and quiescence and are modulated by environmental conditions (including social context) and internal state ^8,9,12,16,18–20^. In the social context, huddling provides thermal and energetic benefits through shared body heat and reduction of heat loss ^21–24^. During mild cold stress, huddling among mice facilitates both entry into, and exit from, an energy-saving, low-Tb quiescent state ^12^. In the non-social context, physical activity and nesting are two behavioral strategies to generate and retain body heat as temperature decreases ^13,25–27^. These patterns suggest active neural regulation of behavior and sympathetic arousal in tandem; however, the brain circuits that support this coordination remain unidentified.

Oxytocin (OT)-producing neurons in the paraventricular nucleus of the hypothalamus (i.e., PVN^OT^) are a compelling candidate for such coordination ^28^. PVN^OT^ neurons have been implicated in autonomic arousal ^29^, energy expenditure ^30^, and thermoregulation—including activation of BAT thermogenesis ^31–33^. The oxytocin system is also classically associated with social behaviors ^34^ including physical touch ^35^—a key element of huddling. Indeed, OT-mutant rat pups display deficits in both warm-seeking and huddling behavior ^36^, and OT neurons display bursts of pulsatile activity during bouts of lactation and direct physical contact with pups ^37^. Yet, whether PVN^OT^ neural activity is functionally associated with the rhythmic patterning of thermoregulatory states is not known.

Here, we set out to identify the relationship between PVN^OT^ activity and daily patterns of behavioral thermoregulation in mice. We initially observed that PVN^OT^ activity is linked to huddling states. Then, we discovered that PVN^OT^ calcium dynamics during huddling are associated with increased likelihood of transitions to body warming and arousal. Intriguingly, in a parallel fashion, PVN^OT^ activity also tracks transitions from rest to arousal in solo animals. Last, using automated thermal feature tracking and optogenetics, we demonstrate that PVN^OT^ stimulation initiates state-dependent transitions that coordinate physiological warming with behavioral arousal. These findings reveal a previously unrecognized role for PVN^OT^ neurons in regulating dynamic, context-dependent thermo-behavioral strategies.

## Results

### PVN^OT^ neural activity is associated with huddling substates

Mice use huddling to facilitate thermoregulatory state transitions. We sought to identify huddling-associated brain regions in groups of adult females, which show stronger huddling-associated Tb changes than male groups ^12^. Using a home-cage surveillance system that eliminates direct human-animal interaction ^12^ in conjunction with FOS immunohistochemistry (IHC), we examined patterns of cellular FOS activity in 18 brain regions. We focused on regions associated with thermoregulation, social behavior, and identified to be FOS-activated during huddling substates in preliminary studies (Fig. 1A-B). We examined animals engaged in active huddling (close physical contact while displaying some physical activity), quiescent huddling (close physical contact with no physical activity) and—as a control for physical touch in the absence of social contact—solo-housed animals engaged in self-grooming. For IHC, we scored cellular FOS activity after the animals had been engaged in state-specific behavior (i.e., active huddle, quiescent huddle, solo-groom) for at least 15 minutes.

**Figure 1.**
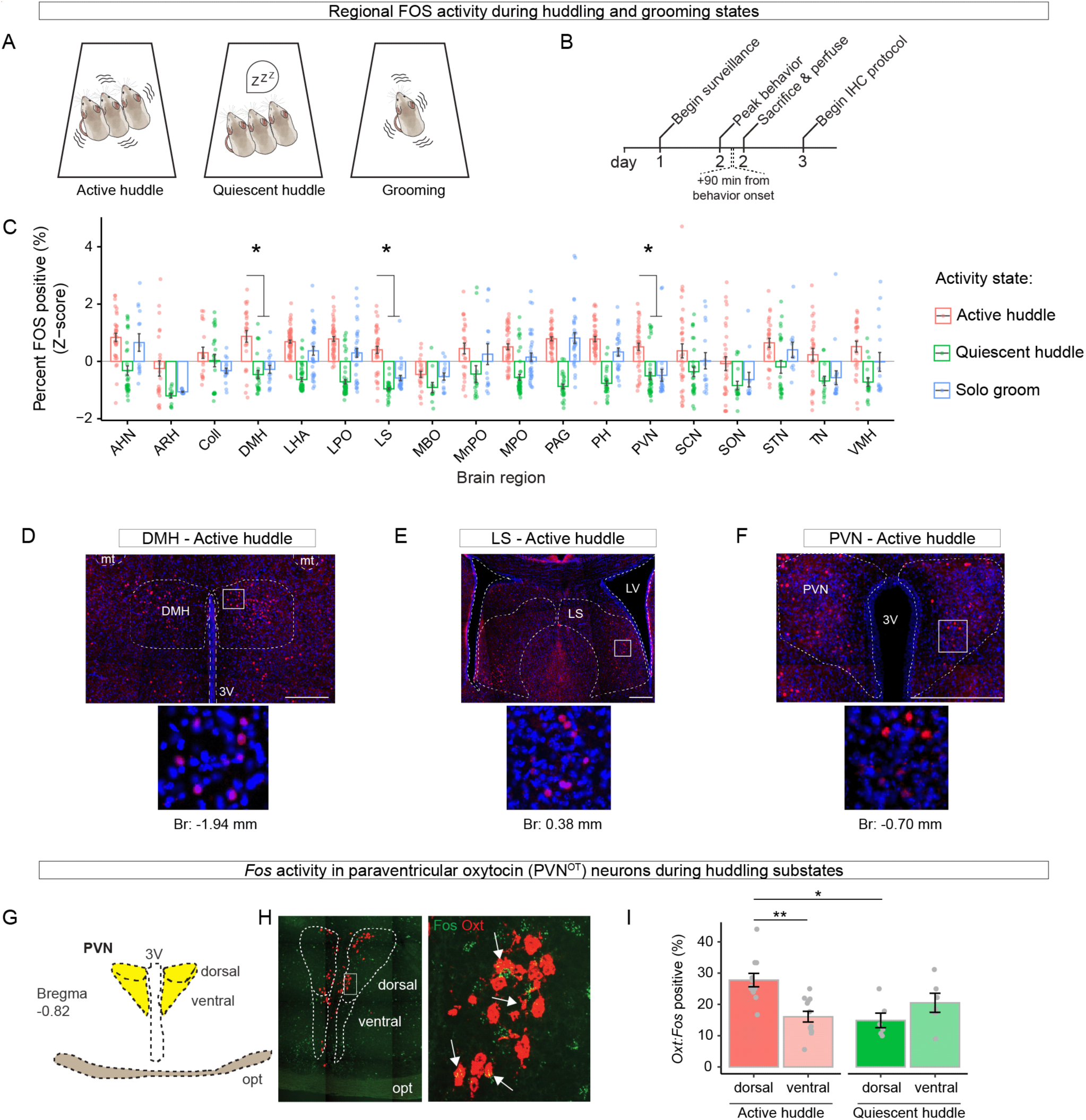
The PVN and PVN^OT^ neurons are FOS activated during huddling substates. **(A - B)** Experimental design to identify state-specific activity in 18 brain regions. Behavioral states **(A)**. Experimental timeline **(B)**. **(C)** Quantification of percent FOS - DAPI colocalized cells across brain regions for each behavior. Regional FOS percentages are z-scored on a per experiment basis. N = 15 mice/1530 ROIs. Each datapoint is an ROI. Asterisks denote regions in which active huddle FOS activity is greater than both quiescent huddle and solo groom. P-values adjusted for multiple comparisons using the Holm method. **(D - F)** Representative histology images showing active huddling associated FOS expression in the DMH **(D)**, LS **(E)**, and PVN **(F)**. 3V: third ventricle; mt: mammillothalamic tract; LV: lateral ventricle. Scale bar equals 500 µm. Insets show FOS (red) and DAPI (blue). **(G)** Schematic of the PVN. Opt: optic tract. **(H)** Representative histology images showing mRNA expression of *Fos* and *Oxt* in the PVN. **(I)** Quantification of percent *Oxytocin:Fos* colocalized cells in dorsal and ventral subregions of the PVN during active huddling and quiescent huddling (N = 8 mice). Each datapoint is an ROI. **C:** linear model with Tukey’s post-hoc tests. **I:** linear mixed effect model. Data are mean ±SEM. P < 0.05 *, P < 0.01 **, P < 0.001 ***. Full statistical analysis in Table S1.

We initially focused on active huddling-associated FOS expression because this state requires that an animal actively maintain close physical contact with another individual. We identified three regions that were activated 90 minutes after the onset of active-huddling: the dorsal medial hypothalamus (DMH), the rostral lateral septum (LS), and the paraventricular nucleus of the hypothalamus (PVN) (Fig. 1C-F and S1A-F; all statistical results are reported in Table S1). The DMH and the PVN are areas associated with energy regulation and thermoregulation, with inputs to the rMR ^5,38,39^. The LS is associated with a variety of functions, from motivation to fear responses ^40,41^ and is strongly associated with the control of social behavior ^42^. These results suggest active huddling recruits brain regions with known roles in thermoeffector and social interaction pathways.

The PVN is one of two primary locations of OT-producing neurons ^39^. Given the established associations of PVN^OT^ neurons with thermoregulation and social behavior, we asked whether they were activated during active and quiescent huddling. To increase the temporal resolution of our assay from the timescale of protein translation to that of mRNA transcription, we used single-molecule fluorescence *in situ* hybridization (smFISH) to identify PVN^OT^ neurons that became *Fos*-positive 30 minutes after the onset of a behavioral state. We quantified neurons in the dorsal and ventral PVN, which have unique cyto-architectures and projection patterns ^43^ (Fig. 1G-I). PVN^OT^ neurons were *Fos*-positive during both active huddling and quiescent huddling. During active huddling, the dorsal population was more *Fos*-positive than the ventral population (*t* = 4.31, *p* = 1.01*X* 10^−3^, LMM, Table S1), and this dorsal population displayed greater *Fos* activity during active compared to quiescent huddling (Fig. 1I) (*t* = 3.30, *p* = 3.94*X* 10^−2^, LMM, Table S1).

These results suggest the PVN is a huddling-associated brain region and that PVN^OT^ neurons are activated during huddling substates. Given the coarse timescale of *Fos* dynamics (≥30 minutes for transcription and ∼60-90 minutes for translation) compared to the faster timescale of rest-arousal cycles in mice (minutes; ^44,45^), we next employed in vivo calcium recordings to capture finescale population-level dynamics of PVN^OT^ neurons.

### PVN^OT^ Ca^2+^ peaks track rest and arousal behavior states in social and non-social conditions

Behaviorally, mice modulate body temperature through physical activity, nesting, and huddling ^7,11–13,25^. In a longitudinal study accounting for individual variance in physiology and behavior, we next asked how calcium activity in PVN^OT^ neurons was associated with real-time behavior during social thermoregulation ^46^ in different thermal conditions (Fig. 2A-C). As a control for social context, we also examined solo-housed animals. Experiments were conducted at three floor temperatures: cold (15**°**C), cool (23**°**C; room temperature), and a temperature near the “lower critical temperature” (29°C; the transition point between energy-requiring and energy-neutral thermoregulatory mechanisms ^11,47,48^). We tracked six behaviors in the solo condition and 10 behaviors in the paired condition (Fig. 2B).

**Figure 2.**
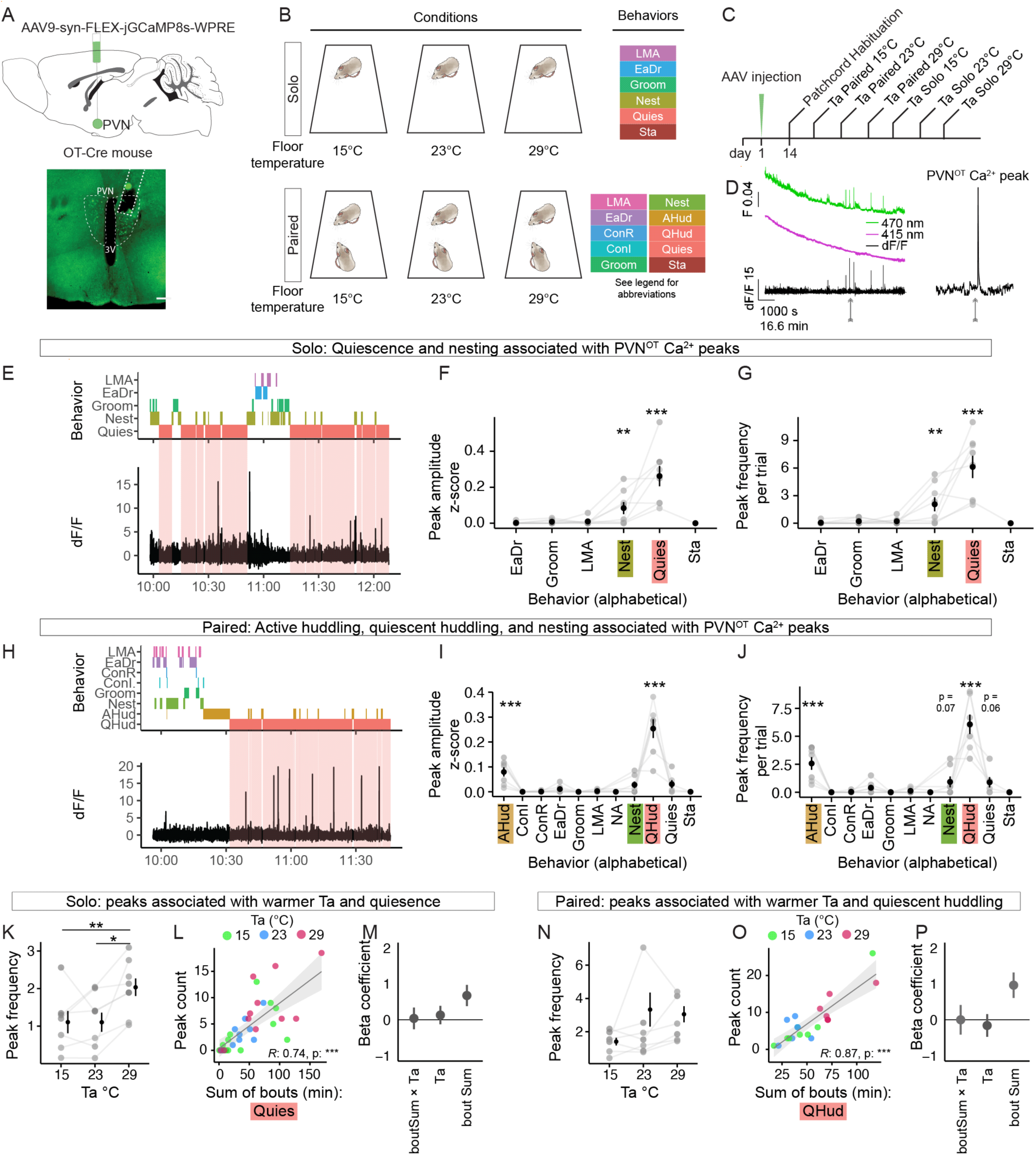
PVN^OT^ peaks track rest and arousal behavior states in social and non-social conditions. **(A - C)** Fiber photometric recordings of PVN^OT^ cells in different social contexts and floor temperature conditions. Scheme of GCaMP AAV injections (top) and histology showing GCaMP-positive neurons in the PVN optic fiber placement (bottom) Scale bar 200 µm. **(A)**. Floor temperature and social context conditions, and behaviors analyzed **(B)**. Scheme of the timeline. Order of the social and floor temperature conditions was pseudo-randomized **(C)**. **(D)** Example traces of calcium-dependent (470 nm) and calcium-independent (415 nm) channels, and the post-processed dF/F trace (bottom), from a two-hour recording. Example PVN^OT^ Ca^2+^ peak, denoted by arrow, is expanded (right) **(E - G)** Behaviors associated with PVN^OT^ peaks in solo condition. Example ethogram aligned to photometric recording. Soft red shaded areas align with the quiescent state **(E)**. PVN^OT^ peak amplitude **(F)** and frequency **(G)** according to behavioral state across solo trials. **(H - J)** Behaviors associated with PVN^OT^ peaks in paired condition. Example ethogram aligned to photometric recording. Soft red shaded areas align with quiescent huddling **(H)**. PVN^OT^ peak amplitude **(I)** and frequency **(J)** according to behavioral state across paired trials. **(K - M)** Effect of floor temperature and quiescence on PVN^OT^ peaks in solo animals. Peak frequency according to floor temperature (Ta) **(K)**. Peak count according to total duration of quiescent bouts **(L)**. Beta coefficients from a model of the effect of floor temperature, total duration of bouts (boutSum), and the Ta*boutSum interaction on peak count **(M)**. **(N - P)** Effect of floor temperature and quiescence huddling on PVN^OT^ peaks in paired animals. PVN^OT^ peak frequency according to floor temperature **(N)**. Peak count according to total duration of quiescent bouts **(O)**. Beta coefficients from a model of the effect of floor temperature, total duration of bouts (boutSum), and the Ta*boutSum interaction on peak count **(P)**. **F,G,I,J,K,L,M,N,O,P**: linear mixed model. N = 8 mice/50 recordings. **M** and **P** show means plus confidence interval; all else shows mean ±SEM. P < 0.05 *, P < 0.01 **, P < 0.001 ***. Full statistical analysis in Table S1. Abbreviations: LMA (Locomotor Activity), EaDr (Eating or Drinking), Groom (Grooming), Nest (Nesting or Nest Building), Quies (Quiescence), Sta (Stationary), ConI (Contact Initiated), ConR (Contact Received), AHud (Active Huddle), QHud (Quiescent Huddle).

We used an AAV encoding the calcium indicator GCaMP (pGP-AAV9-syn-FLEX-jGCaMP8s-WPRE) ^49^ and fiber photometry using an optical fiber implanted above the PVN (Fig. 2A-D). GCaMP AAVs have been validated to transfect OT neurons in OT-Cre mice ^50–52^ and, in this study, showed co-labelling with OT-immunoreactive neurons (Fig. S2A-B). Neural-photometric recordings were conducted during the light/rest phase, when rhythmic episodes of rest and activity make the distinction between different thermoregulatory states readily discernable in female (but not male) mice ^12^. We observed large calcium transients (hereafter “peaks”) that appeared to be similar to those previously reported in lactating female mice ^37,51^ (Fig. 2D). Further analyses focused on peaks that were at least six standard deviations above baseline.

We first examined the effect of social context (i.e., paired- vs. solo-housed) on calcium peak amplitude and frequency across the three floor temperatures. While there was no effect of social context on peak amplitude (Fig. S2C), there was an increase in peak frequency in paired animals (Fig. S2C-D) (*t* = 2.455, *p* = 1.82*X* 10^−2^, LMM, Table S1). To our surprise, in both social and nonsocial contexts, PVN^OT^ peaks were strongly linked to specific behaviors. In the solo context, peak amplitude and frequency were associated with nesting (*t* = 2.83, *p* = 7.64*X* 10^−3^, LMM, Table S1) and quiescence (i.e., motionless rest) (*t* = 8.85, *p* = 1.98*X* 10^−10^, LMM, Table S1) (Fig. 2E-G). In the paired context, peaks were associated with active (*t* = 5.26, *p* = 2.35*X* 10^−6^, LMM) and quiescent huddling (Video S1) (*t* = 16.74, *p* = 1.84*X* 10^−23^, LMM) and, to a lesser extent, nesting (Fig. 2H-J, Table S1). We next examined the behavioral associations of PVN^OT^ peaks within each floor temperature. The behavior-state associations with calcium peaks were largely preserved across all three floor temperatures (Fig. S2G-J). Linear mixed model (LMM) analysis of calcium peak amplitude and frequency revealed main effects of quiescence and nesting in the solo context (Fig. S2G,I), and main effects of active huddling, quiescent huddling, and nesting in the paired context (Fig. S2H,J). Thus, PVN^OT^ peaks are associated with distinct resting and waking behavioral states in social and non-social contexts.

Next, we analyzed the relationship between peak frequency and floor temperature and behavioral bout length across the two social contexts (Fig. 2K-P). For resting behaviors (quiescence and quiescent huddling), peaks were more frequent at 29**°**C for solo (Fig. 2K) animals, and trended (non-significantly) upward in paired animals (Fig. 2N, Table S1). In mice, sleep is more common at warmer ambient temperatures ^8^. To disambiguate whether PVN^OT^ peaks display warm sensitivity vs. arousal-state dependency, we performed multiple regression analysis relating the number of peaks to floor temperature and the duration of rest bouts. In solo and paired conditions calcium peak counts were strongly related to rest bout duration (Fig. 2L,O), and quiescence and quiescent huddling bouts appeared longer at 29**°**C (Fig. 2L,O, red dots). Analysis of the contribution of bout length and floor temperature to peak counts showed that, in solo and paired contexts, increased peak counts at 29**°**C could be explained by longer rest bout lengths (Fig. 2M and 2P). We then examined active behaviors associated with PVN^OT^ peaks. For active huddling, peak counts were positively correlated with bout length, but not temperature (Fig. S2K-L). For nesting, peak counts were not associated with bout length nor floor temperature (Fig. S2M-N).

Together, these results suggest PVN^OT^ peaks track distinct resting and active behaviors in social and non-social settings and are enhanced in the social context. In addition, PVN^OT^ peaks are largely unaffected by the floor temperature conditions tested; we therefore pooled data from the three floor temperatures in subsequent analyses.

### PVN^OT^ neuronal activity precedes transitions towards thermogenesis and behavioral arousal in social and non-social contexts

PVN^OT^ peaks track two rest states (quiescence and quiescent huddling), and two active states (i.e., nesting and active huddling) (Fig. 2). Notably, each of these states are in part thermoregulatory and are naturally organized into alternating sequences of bouts across rest/active cycles ^8,9,12,16,18–20^. To address how PVN^OT^ dynamics align with these sequences, we quantified peaks relative to the onset and offset of behavior bouts, as well as to changes in physical activity and core body temperature (Fig. 3A).

**Figure 3.**
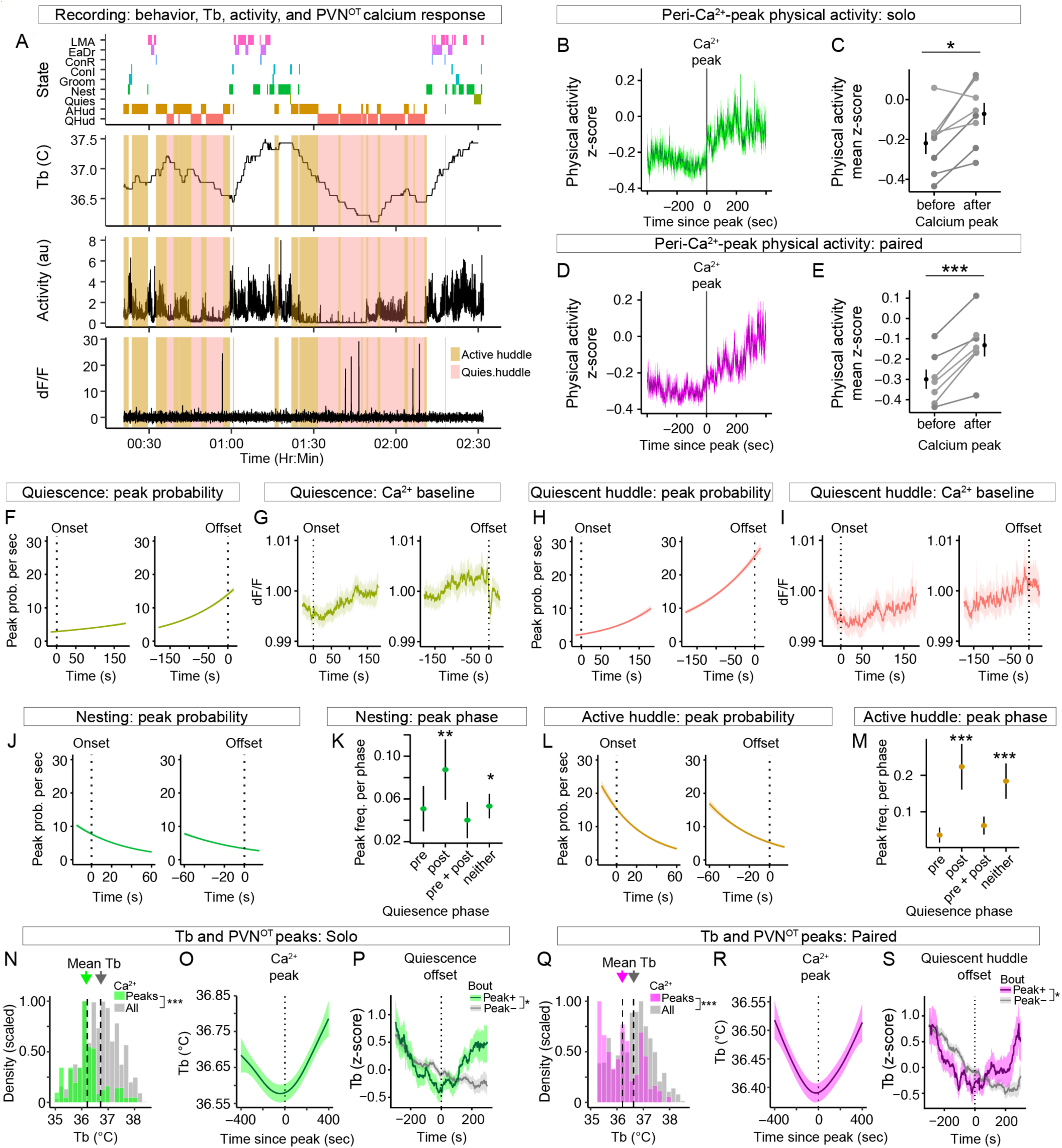
PVN^OT^ peaks are associated with increased likelihood of thermogenic rest-to-active transitions. **(A)** Individual, example trace of alignment of behavioral state, body temperature, physical activity, and dF/F from an experiment in the paired context. Bouts of active- and quiescent-huddle are color coded. **(B - E)** Physical activity around the time of PVN^OT^ peaks. Peri-event time histogram of activity before and after PVN^OT^ peaks in solo mice **(B)**. Per-individual means before and after PVN^OT^ peaks in solo mice **(C)**. Peri-event time histogram of activity before and after peaks in paired mice **(D)**. Per-individual means before and after peaks in paired mice **(E)**. **(F - I)** PVN^OT^ Ca^2+^ dynamics during onset and offset of rest-behavior bouts. Quiescence onset/offset: peak probability **(F)** and Ca^2+^ baseline average **(G)**. Quiescent huddle onset/offset: peak probability **(H)** and Ca^++^ baseline average **(I)**. **(J - M)** PVN^OT^ Ca^2+^ dynamics during onset and offset of active-behavior bouts. Nesting onset/offset: peak probability **(J)**. Nesting-associated peak frequency according to quiescence phase. “Neither” refers to bouts not adjoining bouts of quiescence **(K)**. Active huddle onset/offset: peak probability **(L)**. Active-huddle-associated peak frequency according to quiescence phase. “Neither” refers to bouts not adjoining bouts of quiescence **(M)**. **(N - S)** Relationship between core body temperature (Tb) and PVN^OT^ Ca^++^ dynamics and behavioral transitions. **(N-P)** Tb dynamics in solo animals. Histogram of Tb during minutes containing a calcium peak (green) vs. baseline (grey) data in solo animals **(N)**. Event-triggered average of Tb before and after PVN^OT^ peaks in solo animals **(O)**. Scaled Tb transitions around quiescence offset for bouts with PVN^OT^ peaks (Peak+) and without peaks (Peak-). Event-triggered average of z-scored body temperature aligned to the time of quiescence offset (0 s). Peak+ bouts had a Ca^2+^ peak within 100 s preceding bout offset. Lines show mean z-Tb; shaded ribbons indicate SEM **(P)**. **(Q – S)** Tb dynamics in paired animals (similar to N-P). Histogram of Tb during minutes with a peak vs baseline **(Q)**. Event-triggered average of Tb before and after PVN^OT^ peaks **(R)**. Event-triggered average of Tb aligned to quiescent huddle offset during bouts with and without peaks **(S)**. **C,E,K,M,N,P,Q**: linear mixed model. **F,H,J,L**: logistic regression. N = 8 mice/50 recordings, except for **N-S**, N = 5 mice/24 recordings. **O** and **R** :predicted values of a general additive model ±SEM; **P** and **R**: linear mixed model. All data shows mean ±SEM. P < 0.05 *, P < 0.01 **, P < 0.001 ***. Full statistical analysis in Table S1.

We first examined physical activity (as measured by frame-to-frame pixel changes). Physical activity increased following PVN^OT^ peaks in both solo (*t* = 3.37, *p* = 0.012, LMM) and paired (*t* = 6.82, *p* = 0.001, LMM, Table S1) conditions. This increase began at the approximate time of the peak and persisted through the subsequent 400 seconds (6.6 min) (Fig. 3B-E).

We next examined the density (i.e., frequency of peaks per time bin) of calcium peaks with respect to the onset and offset of the two resting states: quiescence and quiescent huddling. For both behaviors, there was a higher density of peaks in the minutes prior to bout offset compared to onset (Fig. S3A-C). Moreover, there was a positive relationship between bout length and the number of peaks per bout, indicating that longer rest bouts are more likely to have peaks (Fig. S3B,D). To better understand the relationship between PVN^OT^ dynamics and bouts of quiescence and quiescence huddling, we next examined (1) the per-second probability of observing a peak relative to bout onset/offset using logistic regression, and (2) changes in calcium baseline (Fig. 3F-M).

For bouts of quiescence, peak probability was approximately 2.7% near the time of onset and 14.0% near the time of offset (Fig. 3F). For bouts of quiescent huddling, peak probability was approximately 2.1% near the onset and around 25.6% near the time of offset (Fig. 3H). In accordance with the peak density and probability data, baseline calcium in PVN^OT^ neurons was higher in the minutes leading to rest offset compared to onset in all animals (Fig. 3G,I, see also S3M-N). Thus, PVN^OT^ peaks are at least five-fold more likely to occur near the offset of quiescence/quiescent compared to onset, and signal an increase in physical activity—a correlate of behavioral arousal ^53^ and a means of increasing metabolic rate and Tb ^26^.

We next examined PVN^OT^ peak density and probability during the onset and offset of the two active states: nesting and active huddling (Fig. 3J-M, and S3E-H). In contrast to the two resting states, peak density in both states was higher near bout onset compared to offset (Fig. S3E,G). Moreover, there was only a weak relationship between peak count and bout length for both states, indicating that longer bouts are not more likely to have peaks (Fig. S3F,H). Logistic regression analysis of nesting bouts showed that PVN^OT^ peak probability was approximately 7.7% near the time of onset and 3.3% near the time of offset (Fig. 3J). For active huddling bouts, peak probability was approximately 15.3% near the time of onset and 5.2% near the time of offset (Fig. 3L). Thus, for nesting and active huddling, PVN^OT^ peaks are two- to three-fold more likely to occur at bout onset than offset. Notably, within the paired condition, there was a higher peak probability for active huddling compared to nesting (Poisson regression, nesting vs. active huddling peaks counts, P < 0.001), suggesting that social interaction enhances the probability of PVN^OT^ activity.

Because the pre-onset and post-offset periods can be composed of multiple different behaviors, the calcium analyses described above were restricted to the time-window of the behavioral bouts themselves (i.e., between onset and offset) plus a small (30 sec) margin. To provide a broader view of PVN^OT^ activity across the extended peri-event period, we examined baseline calcium for three minutes before and after the onset and offset of each type of behavior (Fig. S3I-L). For the two resting states (quiescence and quiescent huddling), baseline calcium reached a minimum near bout onset and a maximum near offset (Fig. S3I-J), whereas for the two active states (nesting and active huddling) baseline calcium showed the opposite pattern, with the maximum near onset and the minimum near offset (Fig. S3K–L). Thus, despite heterogeneity in the peri-bout periods, PVN^OT^ baseline calcium exhibits consistent state-dependent dynamics around the offset of rest and onset of active behaviors.

Nesting and active huddling often occur immediately before or after (or in between) bouts of quiescence (i.e., the peri-quiescent phase) ^12,16,19,20^. We next examined the quiescence phase for PVN^OT^ peaks associated with nesting and active huddling. PVN^OT^ peaks were more likely to occur during the post-quiescence phase for both nesting (Fig. 3K) (*t* = 2.97, *p* = 0.005, LMM) and active huddling (Fig. 3M) (*t* = 4.08, *p* = 1.58*X* 10^−4^, LMM, Table S1). Notably, although nesting and active huddling are commonly expressed during the pre-quiescent phase, these two behaviors are rare during the post-quiescent phase (Fig. S3O-P). Thus, even though nesting and active huddling are relatively rare during the post-quiescent phase, those are precisely the active state-phases during which PVN^OT^ activity increases. This suggests that PVN^OT^ neurons distinguish nesting/active huddling in the post-quiescent phase from these same behaviors in the pre-quiescent phase.

Together these results suggest that elevated PVN^OT^ activity dynamics precede the offset of two rest states (quiescence and quiescent huddling) by approximately 100 seconds, and the onset of two post-quiescence active states (nesting and active huddling) by around 20 seconds, in solo and paired mice respectively.

### PVN^OT^ peaks occur during low Tb (∼36.2°C) and prior to body warming

In rodents, transitions from rest to arousal are thought to result, in part, from an ultradian (i.e., occurring on cycles < 24 hr), brain-driven switch of the defended body temperature from a rest balance-point (i.e., lower bound) to an awake balance-point (i.e., upper bound) ^2,7^. Because PVN^OT^ calcium peaks aligned with transitions from resting to active states, we examined Tb dynamics in relation to different behavioral states and the timing of PVN^OT^ calcium peaks.

We first measured the relationship between core Tb and behavioral bout length (i.e., duration) using regression (Fig. S3Q-V). For bouts lasting two or more minutes, Tb was not associated with active huddling, eating/drinking, grooming self, or quiescence. In contrast, Tb was negatively correlated with the length of quiescence huddling bouts (Fig. S3V). This finding supports our previous demonstration that quiescent huddling is an energy saving state in mice ^12^.

We then examined Tb dynamics before, during, and after PVN^OT^ calcium peaks (Fig. 3N-S). PVN^OT^ peaks occurred when Tb was around 0.5**°**C lower than baseline (baseline-Tb: 36.7**°**C; peri-peak-Tb: 36.2**°**C) (Fig. 3N,Q) in both solo (*t* = −38.85, *p* < 0.0001, LMM) and paired (*t* = −42.78, *p* < 0.0001, LMM, Table S1) conditions. Moreover, PVN^OT^ peaks aligned with the low point of a U-shaped body temperature profile: on average, Tb decreased before, and increased after, the time of the calcium peak in both solo and paired conditions (Fig. 3O,R). Together, these results suggest that PVN^OT^ peaks occur during a low Tb trough and mark a subsequent rise in Tb.

We next examined rest-to-active transitions, and asked whether the thermogenic trajectory differed depending on whether a PVN^OT^ calcium peak occurred prior to rest offset. We extracted peri-transition temperature traces (±300 s) aligned to the offset of quiescence and quiescent-huddling bouts and classified each transition as Peak+ if it contained one or more PVN^OT^ peaks in the 100 s preceding bout offset, and Peak− otherwise. To control for inter-animal differences in Tb balance point, temperature was z-scored within mouse (Fig. 3P,S). Analysis revealed that transitions preceded by PVN^OT^ peaks exhibited larger post-offset increases in body temperature than transitions lacking peaks for both quiescence (PeakPlus effect: *F*(1, 161) = 4.60, *p* = 0.033) and quiescent huddling (*F*(1, 113) = 5.64, *p* = 0.019, Table S1). Together, these results indicate that although PVN^OT^ peaks are not present before every rest offset, when they do occur it is associated with an enhanced thermogenic rise during the ensuing rest-to-active transition.

### Lactation as a functional validation of PVN^OT^ recordings

We next sought to verify the oxytocinergic identity of recorded GCaMP cells in the PVN using IHC. Unexpectedly, we found variable overlap between oxytocin immunoreactivity and AAV-GCaMP-positive cells. For example, in PVN tissue slices from one individual the percentage of OT^+^,GCaMP^+^ cells ranged from 80% to 69%, while the percentage of GCaMP^+^,OT^+^ cells ranged from 51% to 24% (Fig. S4A-F). Notably, in slices with low overlap between OT and GCaMP markers there was pronounced OT reactivity in the processes lining the third ventricle, particularly the ventral side and the lumen (Fig. S4D). These observations are in accord with reports that OT IHC can reflect release-competent processes, including those contacting or lining the ventricle ^54,55^, and that OT IHC staining between the ventricle and PVN soma can vary by time of day ^56^. We therefore turned to physiology recordings to verify the cellular identity of the GCaMP recorded cells.

PVN^OT^ neurons display characteristic pulsatile activity bursts during lactation and direct contact with pups, with heightened intensity and broadening waveforms during late-stage lactation (PPD 8 to 14) compared to early-stage lactation (PPD 2-7) ^37,51,57^. Whether this type of activity in PVN^OT^ neurons is exclusive to lactation and nursing events is poorly understood ^37,50,58^. To address whether calcium peaks occurring during transitions from quiescence to active states resemble those during lactation, we recorded from virgin females as described above and then allowed them to reproduce and nurse their young (Fig. 4A-C). As documented in other studies ^37,51^, PVN^OT^ neurons displayed peaks that increased in frequency and amplitude from early-stage to late-stage lactation. This observation provides a physiological verification of the oxytocinergic identity of the recorded neurons in our system (Fig. 4D-E and Video S2). PVN^OT^ peaks in virgin females appeared similar in amplitude in early-stage compared to late-stage lactation and appeared narrower overall (Fig. 4F-I).

**Figure 4.**
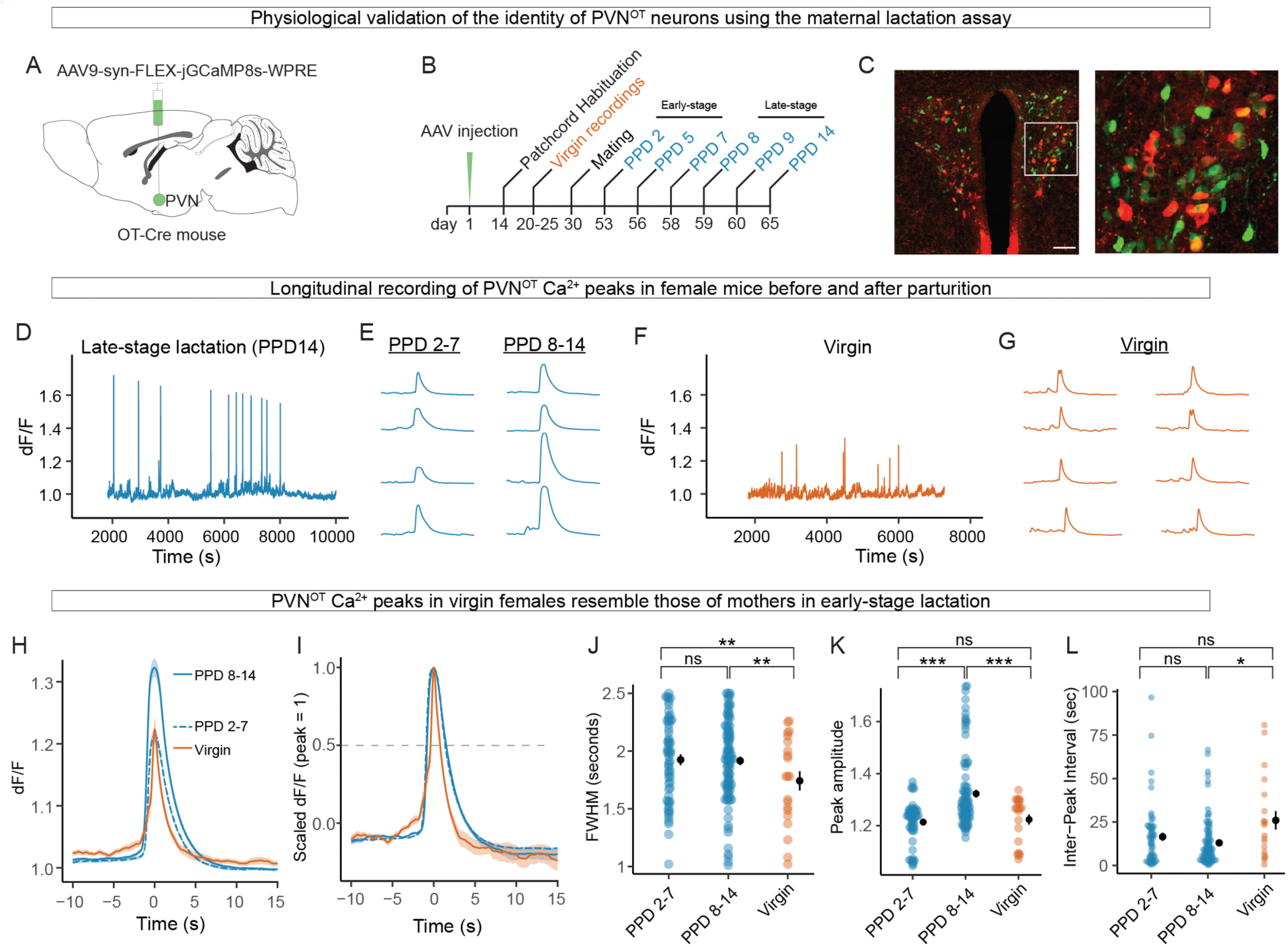
Validation experiment: PVN^OT^ peaks observed in virgin females share similarities with peaks seen in lactating mothers. **(A - C)** Longitudinal fiber photometry recordings of PVN^OT^ cells in virgin-to-lactating mice. Scheme of GCaMP8s AAV injections **(A)** and recording timeline **(B)**. Typical histology section showing PVN stained with anti-OT (red) and GCaMP8s expression (green). Scale bar 100 µm. **(C)**. (D - G) PVN^OT^ Ca^++^ peaks in virgin females and in mothers during early-stage (PPD 2-7) and late-stage (PPD 8-14) lactation. Example of a post-processed dF/F trace from a two-hour recording on PPD 14 **(D)**. Representative peaks from PPD 2-7 and PPD 8-14 **(E).** Example of a post-processed dF/F trace from a two-hour recording of a virgin female **(F)**. Representative peaks **(G)**. **(H - L)** PVN^OT^ Ca^++^ peak kinetics in virgins and lactating mothers. Average dF/F over time of peaks according to condition shown as raw data **(H)** and scaled (peak-normalized with amplitude = 1) **(I)**. Full-width of half maximum (FWHM) **(J)**, peak amplitude **(K)**, and interpeak interval **(L)** according to female condition. N = 2 females, N = 24 recordings, N = 174 peaks. Full statistical analysis in Table S1.

Systematic comparisons of peak kinetics revealed that virgin female calcium peaks displayed narrower full-width half maximum (FWHM) values than either early-stage or late-stage lactation (Fig. 4I-J). Next, while virgin and early-stage lactation peak amplitudes were equivalent, both were less than that of late-stage lactation (Fig. 4K). Finally, while virgin interpeak intervals (IPIs) were equivalent to early-stage lactation, they were longer than those of late-stage lactation (Fig. 4L) (all statistics in Table S1).

Together, these results provide a physiological validation of the oxytocinergic identity of the recorded neurons. They suggest that calcium peaks observed in virgin females during rest-wake behaviors (this study) share similar kinetics to PVN^OT^ peaks during early-stage lactation. As such, this data uncovers a behavioral and physiological context in which PVN^OT^ neurons display calcium burst-like peaks.

### PVN^OT^ peaks signal increases in BAT thermogenesis and changes in regional surface temperature

Our data suggest PVN^OT^ peaks signal transitions towards thermogenesis and arousal as measured by core Tb and physical activity. However, core Tb represents the integration of several different thermoeffector pathways like BAT thermogenesis and peripheral thermal regulation (e.g., cutaneous blood flow. We therefore sought to automate a computational strategy that uses video thermography in freely moving animals to approximate BAT thermogenesis and <u>regional surface temperatures</u> at sub-second resolution—a timescale relevant to sympathetic activity.

We developed a computer vision system called Skeleton-Guided Bodypart Segmentation (SGBS) that uses deep learning to record the surface temperature of three thermal features of an animal: the surfaces over (1) interscapular brown fat (BAT), (2) the rump, and (3) the entire dorsal surface (Fig. 5A). Our pipeline integrates DeepLabCut-based animal tracking ^59^ and Mask R-CNN technologies ^60^ in a Python framework^61^ to process high-resolution thermography (FLIR) videos. An attention layer integrates 10 anatomical skeleton keypoints (e.g., head, nose, shoulders and tail) with a model trained on manual annotations of the three thermal features. Leveraging these keypoints, SGBS can precisely localize and delineate thermal features for mask prediction (Fig. 5B). SGBS detects above-average temperatures in the BAT region and below-average temperatures for the rump region without the need for shaving the back—a manipulation that alters heat loss and can alter behavioral and autonomic thermoregulation ^62^ (Fig. 5C). We compared the training loss (log scale) for SGBS to an unmodified Mask R-CNN trained under identical conditions. The SGBS model converged more rapidly and reached an approximately 100-fold lower final loss than the baseline network, indicating that incorporating DLC-derived keypoint heatmaps enhance segmentation performance in thermographic images (Fig. 5D). We used SGBS to segment BAT, rump, and dorsal surface features on a frame-by-frame basis, extracting the mean temperature for each feature (Fig. 5A and Video S3). SGBS data were synchronized with timestamps and integrated with other data streams, including calcium traces and behavior tracking. For these analyses, solo-housed females were used to eliminate the confounding effects of heat exchange with social partners.

**Figure 5.**
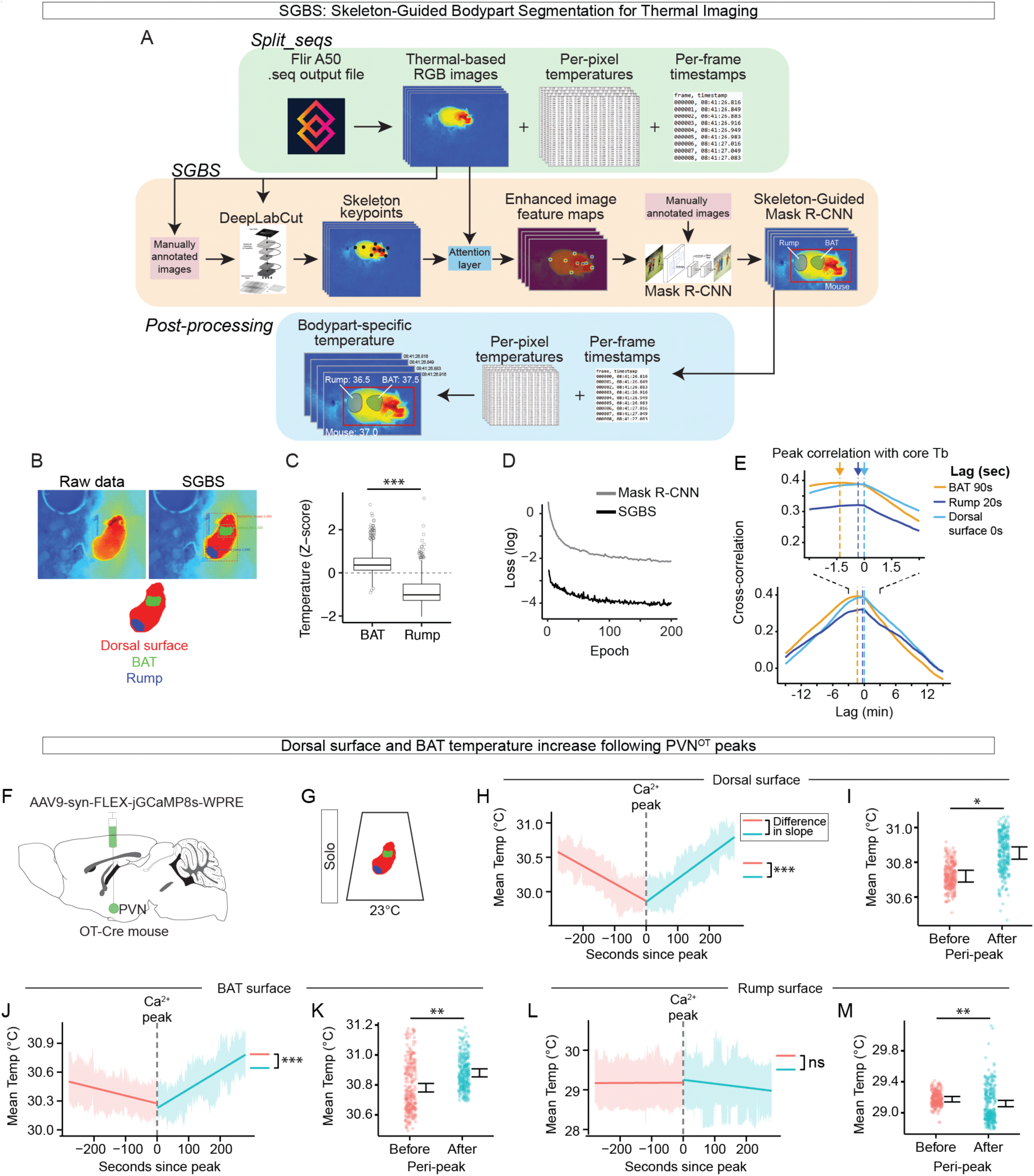
A computer vision model for tracking thermal features during Ca^++^ imaging. **(A)** SGBS model architecture. Three main modules are involved in thermographic identification of surface temperature features: processing raw FLIR .seq files (*Split_seqs*); segmentation of anatomical regions (*SGBS*); overlaying temperatures onto segmented regions for each frame (*Post-processing*). **(B)** Example image showing segmentation of three anatomical thermal features. **(C)** Boxplot shows per-minute mean temperatures (BAT-surface, rump-surface) scaled on a per-mouse basis. **(D)** Loss values between *SGBS* and traditional Mask R-CNN. **(E)** Cross-correlation analysis of core body temperature (Tb) and thermal features (BAT, rump, dorsal surfaces). Data are from light-only control animals recorded for two hours (N = 4). A negative value in the time-shift axis means the thermal features are shifted behind relative to Tb. **(F-G)** Alignment of SGBS thermal feature data with calcium imaging. Viral strategy **(F)** and experimental condition **(G)**. **(H-M)** Peristimulus time histograms (PSTH) of SGBS thermal feature data in relation to the time of PVN^OT^ calcium peaks (time 0). Dorsal surface PSTH slopes **(H)** and means **(I)**. BAT surface PSTH slopes **(J)** and means **(K)**. Rump surface PSTH slopes **(L)** and means **(M)**. N = 4 mice; N = 12 recordings. **H, J, L** slope-lines are fitted values from a linear mixed model. **I, K, M** are per-frame means; statistics from a linear mixed model. **G-M** show means ±SEM. P < 0.05 *, P < 0.01 **, P < 0.001 ***. Full statistical analysis in Table S1.

To test the performance of SGBS, we used thermal recordings from the fiber photometry experiments. We examined the temporal dynamics of pairwise combinations of the three thermal features with respect to core Tb and identified the lag time that gives the best correlation (i.e., cross-correlation analysis) (Fig. 5E). Core Tb and dorsal surface temperatures were very similar with no lag. Next, changes in core Tb lagged changes in BAT by approximately 90 seconds, a result consistent with a previous report ^63^. Finally, rump temperature lagged core Tb by 20 seconds. Thus, SGBS captures the dynamics of thermoregulatory features at the resolution of seconds. These results support the notions that (1) core and dorsal surface temperatures are highly correlated because they both integrate multiple thermoeffector pathways, (2) BAT thermogenesis can drive temperature increases in the mouse body on the order of minutes, and (3) rump surface temperature changes, which may partly reflect vasomotor processes, track Tb on a relatively faster timescale (i.e., seconds) ^64^. Finally, the alignment between the SGBS-derived thermal feature trajectories with independently recorded thermologger core Tb data provides an external physiological validation of the SGBS segmentation pipeline (Fig. 5E).

We next used SGBS to align thermal feature data with PVN^OT^ calcium peaks from fiber photometry recordings (Fig. 5F-M). Consistent with our previous core Tb measurements (Fig. 3N-S), thermal feature temperatures trended downward before the calcium peak but increased afterward for both BAT surface and dorsal surface, a shift confirmed by a significant difference in slope before *vs*. after the peak (Fig. 5H,J) (Table S1). Moreover, BAT (*t* = 2.72, *p* < 0.0065, LMM) and dorsal surface (*t* = 2.48, *p* < 0.0133, LMM) temperatures were significantly greater after the peak compared to before (Fig. 5I,K). By contrast, although rump temperature showed no difference in slope before *vs*. after peak, the mean temperature was lower after the peak compared to before (Fig. 5L-M) (*t* = −3.13, *p* < 0.0018, LMM) (all statistics in Table S1). Together, these results suggest that PVN^OT^ activation is followed with an onset of BAT thermogenesis and a concurrent decrease in rump surface temperature, consistent with sympathetically-driven peripheral heat conservation.

### Optogenetic activation of PVN^OT^ neurons initiates thermogenic responses

To elucidate the functional link between PVN^OT^ activity during quiescence and thermoeffector outputs, channelrhodopsin (AAVDJ-EF1a-DIO-hChR2(H134R)-EYFP-WPRE-pA) was expressed in the PVN of Oxytocin-Cre mice (ChR2+) to allow for optogenetic stimulation of PVN^OT^ neurons (Fig. 6A). To control for potential warm-sensitive physiological responses due to heat from light stimulation, a “light-only” cohort of Oxytocin-Cre mice received optic fiber implants over the PVN. These experiments were conducted at room temperature (23°C) in solo-housed females to focus on the precise physiology of PVN^OT^ neurons and to eliminate confounding effects of social-partners. Blue light stimulation (10 Hz; 20 ms pulse width (20% duty cycle); 10 mW power; 10 or 20 sec pulse train) was delivered after solo animals had been quiescent for at least two minutes (a condition similar to when PVN^OT^ calcium peaks occur). 10 Hz optical stimulation of PVNOT terminals in the rMR elicits thermogenic responses, and 10 Hz stimulation of PVNOT cell bodies increases thermogenesis through a oxytocin receptor dependent fashion in the rMR ^31^.

**Figure 6.**
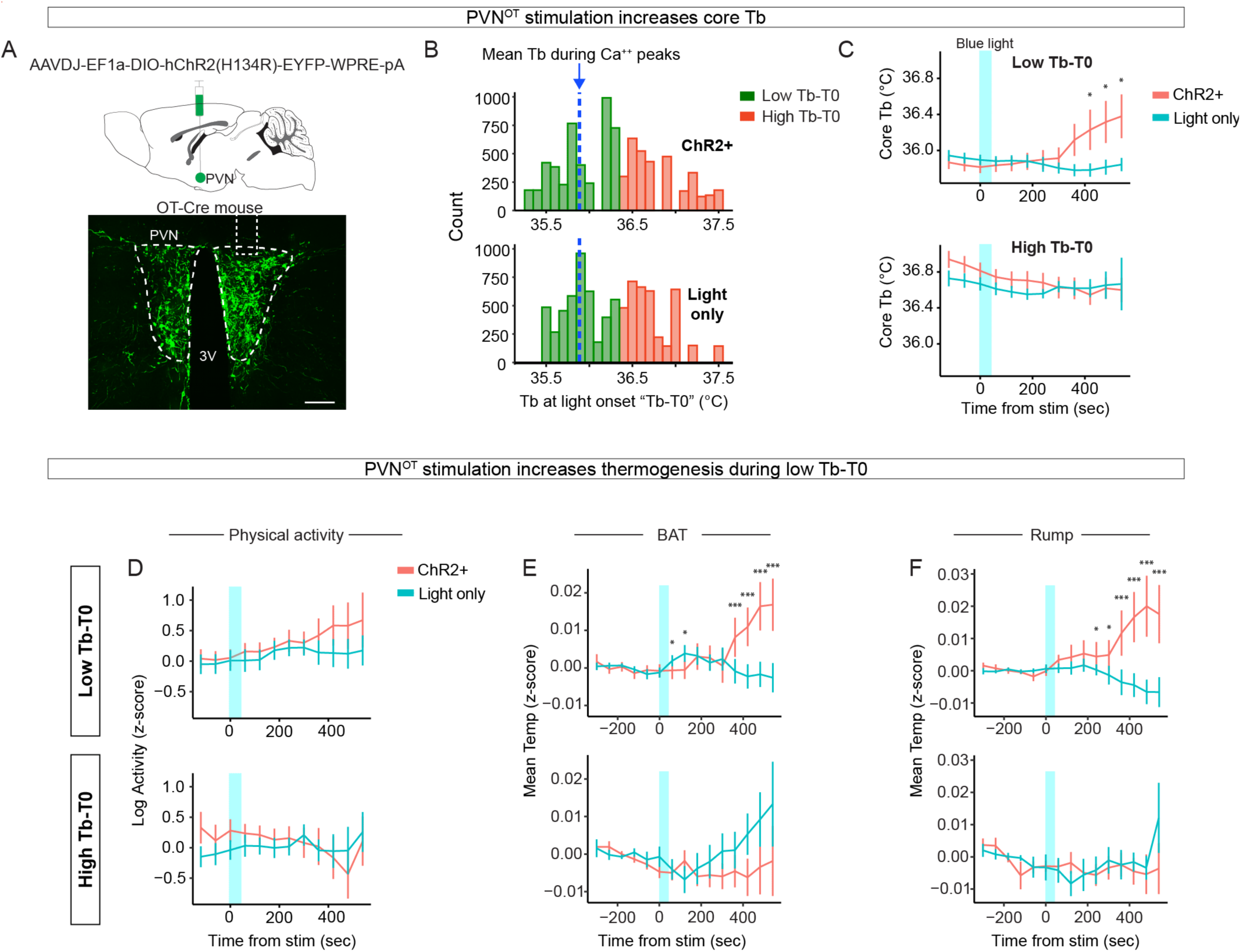
Optogenetic stimulation of PVN^OT^ neurons at a low Tb increases behavioral arousal and thermogenesis. **(A)** Viral strategy for ChR2 transfection. **(B)** Histograms showing core body temperature (Tb) during light stimulations in ChR2+ (top) and light-only controls (bottom). Tb-T0 is the Tb at the onset of light stimulation pulse-trains. Low Tb-T0 (green) refers to stimuli during low Tb, and High Tb-T0 (red) refers to stimuli during high Tb. Blue dotted line denotes the mean Tb at which PVN^OT^ Ca^++^ occur (from Fig. 3N). **(C)** Blue light stimulation (time 0; shaded blue rectangle) increases core Tb during low Tb-T0 in ChR2+ animals compared to light-only animals (top). No effect during high Tb-T0 (bottom). **(D - F)** PVN^OT^ stimulation increases thermogenesis in ChR2+ but not light-only controls. Shown are effects during low Tb-T0 (top row) and high Tb-T0 (bottom row). Optogenetic stimulation (time 0; shaded blue rectangle) effects on physical activity **(D)**, BAT surface temperature **(E)**, and rump surface temperature **(F)** in ChR2+ compared to light-only control animals. **C-F**: linear mixed model on data binned per-minute. Data show means ±SEM. N = 6 mice (3 ChR2+ animals and 3 light-only controls); N = 12 experiments; N = 73 light stimuli. P < 0.05 *, P < 0.01 **, P < 0.001 ***. Full statistical analysis in Table S1.

We first examined the effect of blue light stimulation of PVN^OT^ neurons on core Tb. Approximately 400 sec (6.6 min) after a 20 sec, but not a 10 sec, pulse train, Tb increased in ChR2+ but not light-only controls (Fig. S5A). Further examination of Tb data revealed that, by chance, some blue light stimulations were delivered during a relatively high Tb—in contrast to the low Tb context in which PVN^OT^ calcium peaks naturally occur (Fig. 3N-S). To address a possible mismatch in internal physiology during optogenetic stimulations vs. endogenous calcium peaks, we stratified the data according to whether Tb was above or below average (per-mouse) at the time of stimulation (i.e., Tb-T0). The low Tb-T0 distribution was similar to the Tb distribution observed during PVN^OT^ calcium peaks in solo animals (Fig. 6B blue line, cf. Fig. 3N). We then examined the effect of 20 sec optogenetic stimulation in the stratified data and found ChR2+ stimulation caused increased Tb in the low Tb-T0 context but not the high Tb-T0 context; this increased Tb was observable 420 sec after stimulation (*t* = 2.235, *p* < 0.025, LMM) and persisted at least for several minutes (Fig. 6C) (Table S1). Thus, precisely timed, context-dependent stimulation of PVN^OT^ neurons is sufficient to increase core Tb.

Next, we examined the effect of PVN^OT^ stimulation on physical activity and behavior (Fig. 6D, S5B-C). While no significant differences were detected between ChR2+ and controls, ChR2+ mice showed a trend toward increased physical activity over time in the low Tb-T0 context (Fig. 6D) (Table S1). A separate slope-based analysis of post-stimulation activity revealed that ChR2+ animals exhibited a steeper increase in physical activity compared to light-only controls in the low (but not high) Tb-T0 context. (Fig. S5C). Analysis of manually-annotated behaviors suggested that PVN^OT^ stimulation did not activate a specific motor pattern output but instead resulted in combined increases in the time spent in nesting (LMM estimate coefficient of ChR2+ stimulation: +38.0 sec), locomotion (+54.0 sec), and grooming (+14.5 sec), but not in eating/drinking (−0.4 sec) (Fig. S5B).

We next examined the effect of light stimulation on thermal features using SGBS. Compared to light-only controls, 20 sec pulse trains triggered increased BAT surface (*t* = 3.427, *p* < 6.09×10^−4^, LMM) and rump surface (*t* = 6.921, *p* < 4.48×10^−12^, LMM) temperatures in the low (but not high) Tb-T0 context approximately 360 sec post stimulus (Fig. 6E-F) (all statistics in Table S1). Moreover, in the low Tb-T0 context, blue light stimulation in ChR2+ animals increased the slopes of the surface temperatures of BAT and rump surface temperatures compared to light-only control animals (Fig. S5D-F). In contrast, in the high Tb-T0 context, blue light stimulation did not affect the BAT or rump surface temperatures and caused a slight decrease in the dorsal surface temperature (Fig. S5D-F) (Table S1).

To better understand the temporal ordering of thermogenic pathways with and without optogenetic stimulation, we next turned to two analyses using cross-correlation and lagged regression. In the first analysis, we examined this progression across all mice in the optogenetic cohort, similar to the analysis of the fiber photometry cohort. Here, we computed peak cross-correlation lags between the derivatives of Tb (i.e., dTb/dt), BAT-surface, and physical activity, and found that, in most mice (five out of six, 83%), BAT surface temperature changes preceded changes in dTb/dt, whereas activity did not consistently precede dTb/dt and often followed it (Fig. S5G). Because this dataset is enriched for repeated rest bouts during the inactive phase, this finding is consistent with the idea that during natural rhythms of rest and arousal thermogenesis precedes increases in physical activity ^2,7^.

In a second analysis, we quantified the time course by which optogenetic stimulation influenced core temperature while accounting for concurrent changes in physical activity and BAT surface temperature. We modeled the rate of change in core temperature (dTb/dt) using a mixed-effects lagged regression in which stimulation was represented as a brief input whose influence could persist and decay over 30 seconds. To control for behavioral and thermal contributions, we included activity and BAT surface temperature terms measured at the current time and at preceding time points (0–60 s in 10 s steps). We then compared this to an identical model without the stimulation term. Adding the stimulation term significantly improved model fit (likelihood ratio test, χ² = 7.15, p = 0.0056; Fig. S5H), indicating that stimulation-associated changes in dTb/dt are not fully explained by activity and BAT surface dynamics.

Together, these data suggest that 20 seconds of PVN^OT^ stimulation during the low Tb-T0 resting states is sufficient to trigger a sustained increase in Tb, and to a lesser extent, an increase in physical activity. Thus, PVN^OT^ neurons signal state dependent transitions in thermogenesis and behavioral arousal. These findings are consistent with the notion that natural rhythms of rest to arousal transitions are initiated by a brain-driven switch to a higher defended Tb of the awake state, and that these transitions are often driven by step-wise activation of thermogenic pathways and physical activity ^7^.

### PVN^OT^ cellular projections to the rMR

We next asked what pathways could PVN^OT^ neurons promote transitions towards thermogenesis and physical activity. rMR efferent neurons regulate non-shivering thermogenesis, cutaneous vasoconstriction, and heart rate ^5^, and the surrounding gigantocellular reticular nucleus (GiA) regulates locomotion and movement ^65,66^. In rats, PVN^OT^ to rMR projections (PVN^OT^→rMR) stimulate thermogenesis and cardiovascular responses in an oxytocin receptor (OXTR)-dependent fashion ^31^. To investigate possible PVN^OT^→rMR cell types in mice, we used FluoroGold (FG) to disambiguate magno- vs. parvocellular PVN^OT^ projections ^67^ (Fig. S6A-C). Because the posterior pituitary is located outside the blood-brain barrier, intravenous FG is selectively taken up by terminals of magnocellular neurons and retrogradely transported to their cell bodies. This allows us to infer the neuroanatomical identity (magno- vs. parvicellular) of the PVNOT neurons of interest. The majority of PVN^OT^→rMR projections were non-magnocellular cells located in the caudal and lateral PVN, suggesting they are parvocellular (i.e., PVN^OT(parvo)^) (Fig. S5C), consistent with previous studies ^31,67,68^. In addition, we observed OXTR-positive neurons in the rMR and GiA (i.e., rMR^OXTR^) using smFISH (Fig. S5D). Thus, PVN^OT(Parvo)^ →rMR^OXTR^ neurons are a candidate pathway underlying transitions towards thermogenesis and behavioral arousal in mice.

## Discussion

### State-dependent PVN^OT^ activity during thermo-behavioral transitions

This study identifies a distinct pattern of state-dependent PVN^OT^ activity during natural patterns of thermoregulatory and arousal behaviors. We initially set out to define the role of PVN^OT^ neurons in huddling-associated rest and wake states, but long-duration in vivo recordings revealed large calcium transients (“peaks”) in both social and non-social contexts. These peaks were temporally aligned with transitions from resting states (quiescence and quiescent huddling) to active states (nesting, active huddling, and physical activity), as well as with increases in thermogenesis. Notably, the PVN^OT^ activity patterns and kinetics observed in virgin female mice resembled the well-characterized bursts seen during early lactation ^37,51,69^, suggesting that this pattern of activity has a broader physiological role in coordinating transitions between thermoregulatory and behavioral states. Thus, although oxytocin is often associated with social behavior, our findings indicate that PVN^OT^ neurons provide a state-dependent signal for thermo-behavioral transitions that generalizes beyond the social context.

PVN^OT^ peak dynamics were temporally aligned with the offset of low-Tb resting states. In the minutes preceding the end of a rest bout, baseline calcium increased to a local maximum (Fig. S3I-J) and peaks occurred probabilistically: the per-second probability of a peak rose to approximately 14–25% near rest offset, compared with only ∼2–3% near rest onset (Fig. 3F,H). Logistic regression further supported the conclusion that PVN^OT^activity increases the likelihood of a rest-to-active state transition (Fig. 3F,H,J,L). The presence of peaks that do not immediately precede overt behavior, as seen in the individual trace in Fig. 3A, is consistent with this probabilistic model and suggests that PVN^OT^ activity biases the system toward transition rather than acting as a deterministic trigger.

PVN^OT^ peaks occurred most consistently when core Tb was relatively low, around 36.2°C, approximately 0.5°C below the average rest-phase Tb of C57BL/6 mice in our colony ^12^. In mice, Tb follows ultradian, limit-cycle-like oscillations between biologically constrained lower and upper bounds ^2,7^, and 36.2°C may approximate the lower bound from which the transition toward the active-state upper bound is initiated ^7^. Consistent with this idea, although PVN^OT^ peaks were not present before every rest offset, bouts that did contain peaks showed a steeper thermogenic rise during the ensuing rest-to-active transition. Likewise, both endogenous calcium activity and optogenetic stimulation during a low-Tb (but not high-Tb) rest state signaled increases in Tb, BAT-surface, and locomotor activity within minutes, supporting a state-dependent role for PVN^OT^ neurons in promoting thermogenesis and arousal (Fig. 7).

**Figure 7.**
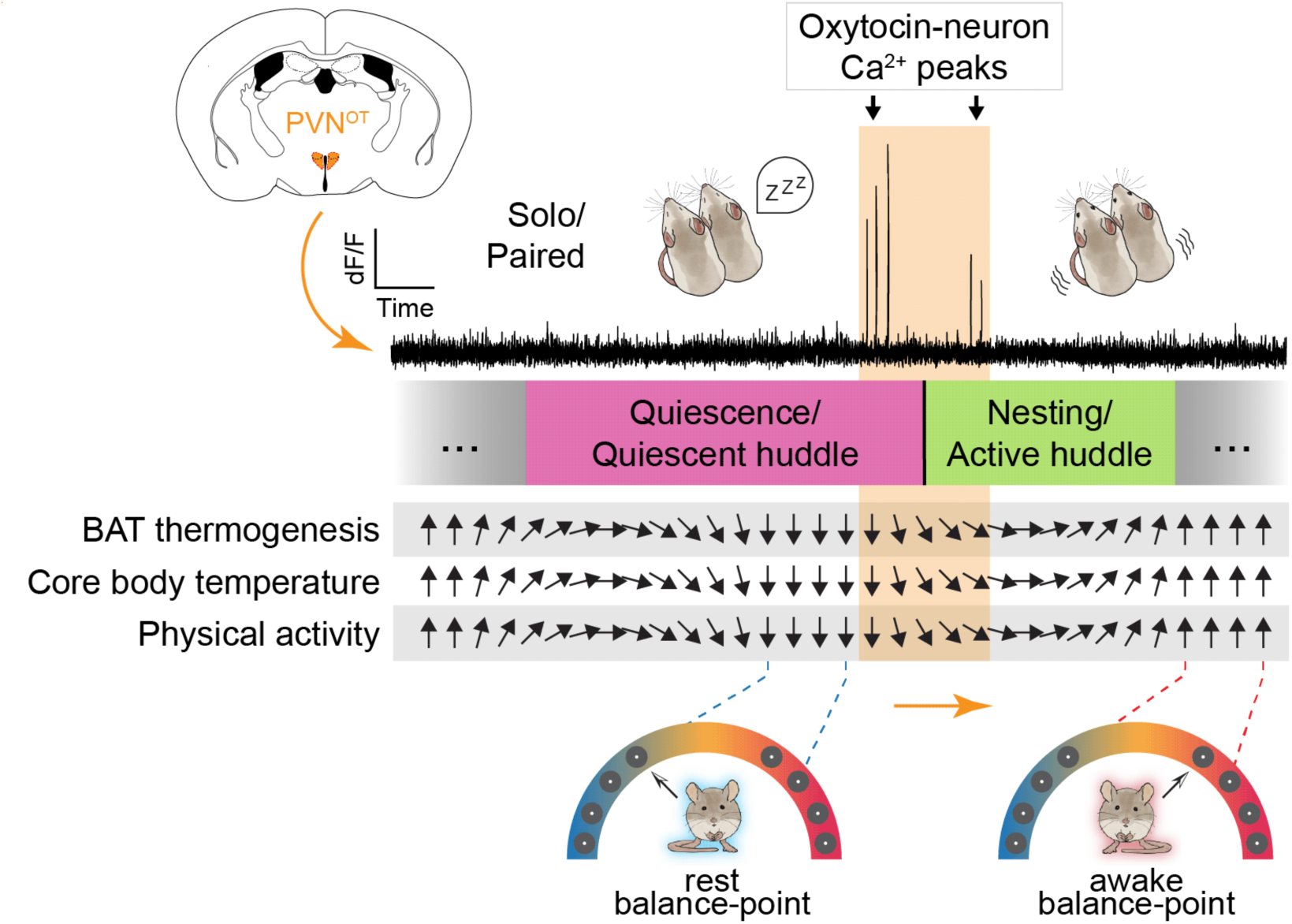
Graphical summary. Increased calcium activity in PVN^OT^ neurons signals the end of rest and the onset of thermogenesis and behavioral arousal. Both the frequency of large amplitude peaks and baseline calcium levels increase approximately two minutes before the end of rest (i.e., quiescence and quiescent huddle). Calcium peaks briefly continue into two post-quiescent active states: nesting and active huddling. Optogenetic stimulation of PVN^OT^ neurons during rest triggers increased BAT thermogenesis, core body temperature (Tb) and physical activity. Arrows indicate transitions between the low-Tb rest balance point (downward vertical arrows) and the high-Tb active balance point (upward vertical arrows).

An apparent contrast emerged between our FOS and fiber photometry results. Whereas FOS expression in the PVN was elevated during active huddling (Fig. 1), photometry showed that PVN^OT^ peaks occurred predominantly during quiescence, immediately before the transition out of rest (Fig. 3). We interpret this difference as a consequence of temporal resolution. Fiber photometry resolves activity on the sub-second timescale and therefore captures transient events occurring while the animal is still behaviorally quiescent. By contrast, FOS integrates neural activity over approximately 30–90 minutes, reflecting cumulative activity across the broader peri-transition period that includes active huddling. Because active huddling is often preceded by quiescent huddling, the associated FOS signals can become intermixed. These findings highlight the complementary strengths of the two approaches and underscore the importance of high-temporal-resolution recordings for identifying neural signals that precede thermo-behavioral state transitions.

The PVN^OT^ activity patterns identified in this study were not specific to social interaction. For example, brief, non-thermoregulatory encounters such as dyadic sniffing did not recruit PVN^OT^ activity, in contrast to the robust peaks observed during transitions out of rest (Fig. S2H; Table S1). However, some aspects of PVN^OT^ activity were enhanced in the social context. For example, peak probability and frequency were higher in the paired compared to solo context (Fig. 3F-I, Fig. S2D). This finding is likely related to our observation that quiescent huddling (paired) bouts were associated with stronger body temperature regulation compared to solo quiescence and other behavioral states (Fig. S3M-N & Q-V). Thus, we uncover a homeostatic function of PVN^OT^ neurons that is social context independent, but nevertheless modulated between the two conditions. Future experiments should attempt to disentangle the effects of PVNOT light stimulation on social vs. non-social aspects of these behavioral state transitions; of particular interest would be to examine how light stimulation affects the duration and thermoregulatory control of social huddling.

### Thermal tracking and validation of PVN^OT^ recording specificity

Our deep-learning pipeline (SGBS) enabled millisecond-resolution tracking of thermal features, including dorsal surface, intrascapular BAT surface, and rump surface temperatures. We detected increases in BAT surface temperature following calcium peaks and optogenetic stimulation, supporting a role for PVN^OT^ neurons in promoting state-dependent transitions toward thermogenesis. Although infrared thermography measures surface radiance rather than tissue temperature directly, the consistent temperature differences between the BAT surface and surrounding dorsal and rump regions (Fig. 5C), together with the temporal lead of BAT surface changes relative to those regions (Fig. 5E and S5G), support the interpretation that the intrascapular BAT region acts as an active heat source during behavioral transitions. While direct telemetry or implanted thermocouples would provide greater precision, our noninvasive approach allowed us to study natural state transitions in freely moving, undisturbed animals. A limitation of the current SGBS system is that it was not designed to track tail temperature, a commonly used index of vasomotor responses.

The lactation model (Fig. 4) served as a functional validation of recording specificity. Although GCaMP expression was driven by an OT-Cre line, co-localization with OT immunoreactivity varied across brain slices (Fig. S4), consistent with prior reports using OT-Cre lines ^70^. We note that the animals were perfused at ∼ZT4–8, before we were aware that somatic OT immunoreactivity in PVN neurons reaches a daily low during the early light phase ^56^. Together with the fact that oxytocin is released from soma and dendrites and must be resynthesized rather than locally recycled ^71^, this timing may have reduced detectable intracellular peptide levels, leading us to underestimate the full OT-expressing population. In situ hybridization targeting oxytocin mRNA, rather than immunohistochemistry, may provide a more reliable assessment of expression specificity. Nevertheless, PVN^OT^ neurons undergo well-characterized changes in firing dynamics during lactation to support milk ejection ^37,51,69^, and the emergence of these characteristic changes in our recorded population (Fig. 4I-L) provides functional evidence that we were primarily recording oxytocin neurons.

### PVN^OT^ neurons in context of arousal and peptidergic PVN cell-types

Several neural populations that promote transitions from sleep to wake have been described ^72^, including populations directly involved in thermoregulation ^9^. Together with a prior report showing that chemogenetic activation of PVN^OT^ neurons increases wakefulness ^29^, our data place PVN^OT^ neurons as a key node in the initiation of thermogenic and behavioral arousal from rest. We therefore propose a model, consistent with prior work, in which the transition from rest to arousal begins with a change in neural activity that drives thermogenesis and is either coincident with, or followed by, increased physical activity ^7,63^ (Fig. 7). The ∼400 s latency between optogenetic PVN^OT^ stimulation and the rise in Tb (Fig. 6) is consistent with the timescale of peptidergic neuromodulation, as oxytocin is released from large dense-core vesicles and can exert prolonged, diffuse effects ^71,73,74^. Our decision to perform optogenetic stimulation in isolated animals was motivated both by the social-context independence of PVN^OT^ activity during rest-to-arousal transitions (Figs. 2 and 3) and by the technical limitation that close physical contact between huddling mice prevents accurate SGBS segmentation of individual thermal profiles.

Other peptidergic PVN populations, particularly CRF and AVP neurons, are unlikely to account for the activity pattern described here. Although their roles in huddling have not been directly tested, available evidence distinguishes both populations from PVN^OT^ neurons. CRF signaling appears to oppose oxytocin in social-care contexts, as chemogenetic activation of PVN^CRF^ neurons impairs maternal care ^75^, and central CRF administration suppresses maternal care ^76^. AVP likewise shows separable functions: intracerebroventricular AVP inhibited nest building but not huddling in *Peromyscus* ^77^, and PVN^AVP^ neurons promote wakefulness via lateral hypothalamic orexin neurons ^78^. Consistent with our findings, Adahman et al. ^79^ also reported reduced PVN^OT^ c-Fos at thermoneutrality relative to cooler conditions in a study of AVP-dependent maternal thermoregulatory behavior. Importantly, chemogenetic activation of PVN^AVP^ neurons did not recruit PVN^OT^ neurons ^80^, indicating these populations can operate independently. The synchronized, pulsatile bursting that validated our calcium peaks appears specific to oxytocin neurons: lactation-induced burst upregulation occurs in magnocellular OT neurons, but not AVP neurons, in the PVN ^81^, whereas AVP neurons exhibit asynchronous phasic bursting associated with antidiuretic rather than neuroendocrine-pulsatile functions ^82,83^. To our knowledge, comparable synchronized burst activity has not been reported in PVN CRF neurons during lactation or rest.

### Limitations and caveats

Caveats of this study should be considered. First, we did not directly link in vivo PVN^OT^ calcium activity or optogenetic stimulation to endocrine oxytocin release. Throughout, we refer to “PVN^OT^ neurons” rather than “oxytocin” to emphasize that our manipulations targeted the cell type, not the neuropeptide itself. PVN^OT^ neurons can co-release glutamate ^84^, and stimulation parameters may influence the recruitment of glutamatergic transmission versus peptide release. High-frequency pulsatile activity is associated with oxytocin release during lactation ^85,86^, whereas lower-frequency stimulation can engage glutamatergic transmission and may interact with oxytocin-receptor-dependent modulation of downstream synapses ^87^. Our optogenetic protocol was nevertheless grounded in rat research showing that 10 Hz stimulation of PVN^OT^ and their projections to rMR can evoke thermogenic responses, including oxytocin-receptor-dependent responses in the rMR ^31^. So while our data show that PVN^OT^ activation is sufficient to initiate state-dependent thermogenic and behavioral transitions, they do not establish whether these effects are mediated by oxytocin, glutamate, or both. That said, we provide anatomical evidence that is compatible with a peptidergic mechanism, including parvocellular PVN^OT^ projections to the rMR along with oxytocin receptor mRNA expression in that region (Fig. S6). Dissecting the relative contributions of oxytocin and glutamate is a next step, requiring receptor-targeted pharmacology or genetic loss-of-function approaches.

Second, although some experiments were conducted in relatively few animals, we nevertheless observed robust within-animal associations between PVN^OT^ activity and thermo-behavioral state changes; future studies with larger cohorts should better resolve inter-individual and state-dependent variation. We focused on females for a practical reason: during the light and rest phase, females show clear, rhythmic bouts of rest and activity, which makes transitions between thermoregulatory states readily discernible and well suited to the analyses around each state transition used here ^12^. This choice constrains the generality of our conclusions, which pertain specifically to females. Because oxytocin signaling can differ between sexes ^88^, and because the neural control of thermoregulatory behavior may not be identical in males, whether the PVNOT dynamics we describe operate similarly in males remains an open question. Testing males directly was beyond the scope of the present study, but it is an important next step, particularly as the neural basis of collective thermoregulatory huddling has recently begun to be defined ^89^.

Third, although we show that PVNOT neurons are sufficient to drive thermogenic and behavioral transitions (Fig. 6), we did not perform acute loss-of-function experiments. Such experiments are warranted because decreases in baseline PVNOT calcium activity were associated with transitions toward the onset of quiescence (Fig. 3I-L), suggesting this system may bidirectionally regulate thermo-behavioral state. A tractable next step would be to optogenetically inhibit PVNOT neurons during established rest bouts, delivered at pseudo random times across the light and rest phase and analyzed post hoc by behavioral state and body temperature; we predict that silencing during rest would prolong the average duration of rest bouts and delay the onset of activity and thermogenesis. Pairing the inhibition with selective oxytocin antagonist (such as L-368,899), would further test whether the thermogenic and autonomic components of these transitions are oxytocin receptor dependent rather than driven by glutamate released by the same neurons.

Finally, our thermal manipulations relied on conductive floor temperature changes rather than whole-chamber ambient air temperature. While floor cooling provided a reliable and well-controlled stimulus that elicited thermoregulatory behavior without altering airflow or humidity, it may recruit peripheral thermoreceptor populations differently than ambient temperature manipulations. Future studies using chamber-wide temperature control would therefore provide an important complement.

### Ideas and Speculation

How might PVN^OT^ neurons detect or define the “rest balance-point,” and contribute to shifting the brain toward an “awake balance-point”? We envision a circuit model in which PVN^OT^ neurons integrate afferent thermosensory information from both peripheral and internal sources to monitor when core Tb approaches a critically low threshold during rest cycles. The lateral parabrachial nucleus (LPB), which serves as a major relay for ascending cutaneous thermosensory information ^5^, is a likely source of peripheral thermal input to the PVN. The preoptic area (POA) contains thermosensitive neurons that monitor brain temperature ^5^ and is poised to relay information to the PVN^OT^ neurons, which receive monosynaptic input ^90^. When Tb drops towards the lower bound of the resting-state thermal range, peripheral and central thermal input may trigger the characteristic pulsatile pattern of PVN^OT^ activity. This activity would then engage downstream effectors, including the rostral medullary raphe, where we detect projections and oxytocin receptor mRNA expression, to initiate BAT thermogenesis and behavioral arousal. Notably, whole-brain projectome mapping has revealed that PVN^OT^ neurons send dense projections to both the parabrachial nucleus and medial preoptic area ^90^. This suggests that PVN^OT^ neurons are not just receiving thermosensory information, but may also modulate LPB and POA in a state-dependent manner through reciprocal feedback.

### Broader Implications

Overall, these findings extend the role of in vivo PVN^OT^ neural activity dynamics beyond social behavior ^35,50,91,92^, parturition/lactation ^58^, maternal behavior ^93^, sexual behavior ^74^, and stress responses ^52^, and highlight their contribution to daily patterns of rest/wake transitions, thermoregulation, and energy balance. In our model PVN^OT^ neurons respond to internal physiological conditions, such as core body temperature, to coordinate autonomic and behavioral outputs necessary for adaptive thermoregulation. These insights open new directions for understanding the neural circuits that integrate social context, internal state, and metabolic demands to regulate mammalian behavior.

## Acknowledgements and Funding

We thank University of Wyoming Sensory Biology Center for input; Robert Carrol for assistance with animal husbandry; members of the Nelson and Bedford laboratory for input; Sean Harrington for coding assistance; the UW Engineering Shop for designing and building hardware. This work is funded by NIH COBRE Grant 5P20GM121310-07.

## Resource availability

### Lead contact

For further information or requests, contact should be directed towards the lead contact, Adam Nelson.

### Materials availability

This study did not generate any new reagents.

### Data and code availability

Data and code underlying this study are publicly available via the Open Science Framework (OSF) at https://osf.io/x9kcj. The repository includes the experimental datasets associated with the main fiber photometry experiments, lactation study, optogenetic experiments, and body temperature–linked photometry experiments, together with metadata files describing session- and animal-level information. Code used for data processing and analysis is also available at this repository.

## Author contributions

M.V., J.G.L, J.F.R., N.L.B., and A.C.N. designed the study. M.V. performed the fiber photometry studies. M.V, J.G.L, and A.C.N. performed the optogenetic studies. M.V., J.G.L., B.A., C.P., and A.C.N. performed the histology studies. J.G.L, S.K. and G.J.T. performed code development. J.F.R. contributed to a previous draft of this manuscript. N.L.B. and A.C.N secured funding and provided expertise and feedback.

## METHODS

### Animals

**OT and FOS histology.** C57B6/J (Strain #: 000664) female mice were purchased from Jackson Laboratory and were 8 to 11 weeks old during the experimental period. Virgin, group-housed females were used for all trials unless otherwise specified.

**PVN^OT^ manipulation and recording.** Oxytocin-Ires-Cre mice were purchased from Jackson Laboratory (Strain #: 024234) (Wu et al., 2012). Animals were bred and maintained at the University of Wyoming. Experimental female mice ranged from 9-20 weeks old during the experimental period. All experimental mice were group-housed with *ad libitum* access to food and water in a room maintained at 23±2°C with a standard 12 h light / dark cycle (light: 6:00-18:00; dark: 18:00-6:00) until they were used for surgery or experiments. Animals were housed in Optimice® IVC cage systems; cages were enriched with red huts, nestlets, and a mix of cob and pine bedding. Mice used were 18-26g during the experimental period. Animal genotyping was outsourced to Transnetyx using ear tissue biopsies. All procedures were approved by the University of Wyoming Institute Animal Care and Use Committee (IACUC) in compliance with NIH guidelines for the care and use of laboratory animals.

### FOS/*Fos* behavior assays

To measure FOS protein expression across brain regions, mice were placed into the home cage recording suite as described (Landen et al., 2024) for at least 24 hours prior to behavior being monitored, with the cage’s camera being connected to a Zoom meeting. After 24 hours, and at the beginning of the light cycle when huddling is most frequent, a human observer monitored behavior from another room without interrupting natural behavior. Once the target behavior was observed for a minimum of 15 consecutive minutes, 90 minutes were counted from the onset of the behavior, and animals were pulled from the recording suite for histology. Animals were not excluded if they exhibited both active and quiescent huddling during the recording session.

To determine *Fos* transcription in PVN^OT^ neurons during different huddling states, C57B6/J female mice that were 10 weeks old were housed in home cages in trios. Mice were habituated to the conditions, handling, and the cage lid camera set up for ∼5 days prior to recording. Animals were split into two groups: active huddle (N=6) and quiescent huddle (N=3). To capture active huddle *Fos* transcription, mice were recorded using the cage lid camera connected to a Zoom meeting. The time of active huddle onset was documented, and mice were sacrificed 30 minutes later. To determine quiescent huddle *Fos* transcription, mice were recorded as described above. Mice were sacrificed 30 minutes after the onset of quiescent or active huddle. All brains were dissected, embedded in OCT (Fisher Scientific, #23-730-571) and stored in −80°C.

### Animal Surgery

**Pre- and post-surgical procedures.** Prior to surgery, animals were given an analgesic dose of carprofen (20 mg/kg, s.c.). Mice were deeply anesthetized using a mixture of ketamine and xylazine (100 mg/kg and 10 mg/kg, respectively, administered i.p.). Post-surgery, the mice were returned to a warm and clean home cage. For post-operative care, antibiotic ointment (Neomycin and Polymyxin B Sulfates and Dexamethasone Ophthalmic Ointment, Bausch + Lomb) was topically applied to the incision, animals received 20 mL/Kg of lactated ringer’s solution s.c., and closely observed until active. Upon emergence from anesthesia, 1 mg/kg of buprenorphine extended-release solution was administered subcutaneously between the shoulder blades. Animals were housed with cagemates and monitored daily. Carprofen was administered for three days post-surgery.

**Temperature logger implants.** To continuously measure internal body temperature, laparotomies were performed to implant mice with Star-Oddi DST nano-T temperature loggers. The loggers were sterilized with glutaraldehyde and rinsed with sterile PBS prior to being implanted. Mice received pre-operative care and were anesthetized as described above. A one-centimeter incision was made in the skin and underlying peritoneum, and the temperature logger was placed into the abdominal cavity. The peritoneum was sutured with PGA absorbable sutures, while the skin was closed with nylon (nonabsorbable) sutures. Mice received post-surgical care as described above.

**Stereotaxic GCaMP injections and fiber implants.** For fiber photometry recordings, oxytocin-Ires-Cre heterozygous, pair-housed, female mice at 8-12 weeks of age underwent stereotaxic surgery in which pGP-AAV9-syn-FLEX-jGCaMP8s-WPRE (#162377, Addgene, titer: 2.7 x 10^13^ gc/mL) was delivered to the PVN. Mice were head-fixed on a digital mouse stereotax (Stoelting, 51730D). For added analgesia, a local anesthetic (5% lidocaine) was delivered subcutaneously along the incision site before cutting. A midline incision was made along the top of the skull, the periosteum was cleared with 3% hydrogen peroxide, and the skull was dried off with canned air. Vertical measurements of bregma and lambda were taken and aligned to the same plane (DV of ± 0.05 mm). The AAV was delivered using a programmable nanoliter injector, (Nanoject III, Drummond Scientific) (300nL per side: 6 cycles of 50nL delivered at 2nL/s with 10 second delays). To allow for proper diffusion of solution into brain tissue, the needle was left in for 10 minutes before retracting. PVN coordinates relative to bregma and DV measured from the top of the brain: AP = −0.76mm, ML= ±0.19mm, DV= − 4.62mm.

A fiber-optic cannula (Ø1.25mm ferrule, Ø200 µm core, NA = 0.37, L= 6.5 mm, Neurophotometrics Ltd.) was then lowered into the injection site until the tip was located ∼ 0.02 mm above the viral injection. The cannula was anchored in place with dental cement (C&B Metabond) such that a dome-shaped cap was halfway up the black ceramic ferrule and covered the entire incision. Post-operative care and recovery took place as described above. Two weeks post recovery, animals were tested for GCaMP signal using the Neurophotometrics photometry system (as described in “**<u>Fiber photometry recordings”</u>**). If a signal was detected, animals underwent a laparotomy to implant a temperature logger as described above. Animals with no detectable signal were excluded from recordings.

**ChR2 injection & optic fiber implant.** For optogenetic stimulation of PVN^OT^ neurons, Oxytocin-Ires-Cre heterozygous, pair-housed, female mice at 8-12 weeks of age were split into two different groups: ChR2+ and control (ChR2-) animals. ChR2+ animals underwent stereotaxic injections with AAVDJ-EF1a-DIO-hChR2(H134R)-EYFP-WPRE-pA (Salk, titer: 4.03×10^13^ gc/mL) using the stereotaxic surgical procedure described above. Viral vectors were delivered using a programmable nanoliter injector, (Nanoject III, Drummond Scientific) (200nL per side: 4 cycles of 50nL delivered at 1nL/s with 10 second delays). Both ChR2+ and control animals were implanted with a fiber optic cannula (Ø1.25mm ferrule, Ø200 µm core, NA = 0.37, L= 6.5 mm, Neurophotometrics Ltd.). All optogenetic animals were given a minimum of seven days to recover before undergoing a laparotomy to implant a temperature logger.

**Stereotaxic DREADD, CTB and FluoroGold injections.** To identify PVN^OT^ projections to rMR, a subset of mice received two stereotaxic injections: one of AAV5-hSyn-DIO-hM3D(Gq)-mCherry (#44361, Addgene, titer: 2.6 x 10^13^ gc/mL) in the PVN and one of cholera toxin subunit B conjugated with Alexa Fluor 647 (Invitrogen, C34778). Using the same surgical procedure and PVN coordinates as mentioned above, 300 nL of AAV was delivered per side (6 cycles of 50nL delivered at 2nL/s with 10 second delays). The incision was closed using absorbable PGA suture threads and antibiotic ointment was applied topically along the incision. Animals recovered for two weeks to maximize viral expression. A second stereotaxic surgery was performed to deliver a volume of 10 nL of 0.5% CTB into the rMR. RMR coordinates relative to bregma and DV measured from the top of the brain: AP = − 5.80mm, ML= 0.00mm, DV= −4.95mm. Mice received post-operative care as described above and recovered for 3-4 days. To distinguish between peripheral-projecting magnocellular and central-projecting parvicellular neurons, mice received 15 uL intravenous injection of 4% Fluoro-Gold (Fluorochrome) diluted in 100 uL of sterile saline. Prior to injection, mice were given an analgesic dose of carprofen (20 mg/kg, s.c.). Mice were briefly restrained using a modified 50 mL conical tube, in which holes were drilled to allow for proper air flow and respiration. Mouse tails were interposed between two heating pads to enhance visibility of the tail vein. Tails were wiped down with 70% ethanol and FG was administered via either right or left lateral tail vein using a 0.5 mL 28G syringe. Mice were sacrificed 24- 48 hours post-FG administration.

### Histology

**FOS Immunohistochemistry.** For quantitative analysis of FOS expression, mice were given a lethal dose of isoflurane after removal from the home cage recording suite. Animals were then transcardially perfused with 1X PBS and 4% paraformaldehyde (PFA) in PBS. Brains were removed and kept in 4% PFA overnight at 4°C. Brains were sliced into 60 um sections using a vibratome (Leica VT1000S) and every other slice was saved in 48-well plates, wrapped in parafilm in 1X PBS, overnight at 4°C. The next day, 0.3 mL of block solution (0.20% triton, 10% natural goat serum (NGS) (Jackson ImmunoResearch 005-000-121), in 1X PBS) was placed into each well and incubated for one hour at room temperature on a plate shaker. Block was removed and replaced with freshly made primary antibody solution (c-Fos (9F6) Rabbit mAb, Cell Signaling, 14609, 1:1000 dilution in block solution) and incubated for 48 hours on a shaker at 4°C. Slices were washed three times with 1X PBS for 15 minutes. Slices were then incubated in secondary antibody solution (Goat anti-Rabbit IgG, Alexa Fluor Plus 555, Thermo Scientific, A32732, 1:500 dilution in block) for 2 hours on a shaker at room temperature and protected from light. Wash steps were repeated with 1X PBS. Slices were then mounted on Superfrost Plus slides using a paintbrush and stained with DAPI mounting medium (Vectashield H-2000). Slides were imaged with a Zeiss Axio Scan.Z1 fluorescent microscope. Regions were identified by finding all locations that displayed FOS expression and cropped using ImageJ alongside the Allen Brain Atlas. The quantity of cells and overlap was calculated using a custom CellProfiler 4.2.1 script.

**Fluorescent *in situ* hybridization (RNAscope).** For quantitative analysis of *Oxt* and *Fos* overlap in PVN^OT^ mice were anesthetized with a lethal amount of isoflurane. Brains were extracted, embedded in OCT and stored at −80°C. Brains were sectioned at 16 µm using a cryostat and subsequently stored at −80°C. For *in situ, s*lides containing brain sections were placed in 4% PFA for 45 minutes and then were dehydrated in 50%, 70% and 100% ethanol for 5 minutes each at RT. Slides dried for 5 minutes after dehydration and hydrogen peroxide was added for 10 minutes at RT. Slides were then treated with Protease III for 30 minutes at RT to allow for antigen accessibility. The probe cocktail contained *Oxt* (ACD Bio, #493171) and *Fos* (ACD Bio, #316921) and was diluted at 1:2 ratio with probe diluent. An aliquot of 40µL of probe cocktail was added to each section and incubated in a 40°C oven for two hours. For amplification, 40µL of AMP 1 was added each slice and placed in the 40°C oven for 15 minutes. AMP 2 was then added to each slice and slides were again incubated at 40°C for 15 minutes. Finally, 2-3 drops of AMP 3 were added to each slice and slides were incubated at 40°C for 30 minutes.

Fluorophores were added to each channel and diluted at 1:1500. Each fluorophore was added by first opening the channel using 2-3 drops of HRP on each slice and incubating at 40°C for 15 minutes. The fluorophore was added at a volume of 40 µL to each slice and slides were incubated at 40°C for 30 minutes. To close the channel, 2-3 drops of blocker were added to each slice and incubated for 15 minutes. Lastly, a drop of DAPI (ACD Bio, #320858) was added to each slice for 30-45 seconds. DAPI was allowed to dry for approximately 3 minutes before immediately coverslipping with 1-2 drops of ProLong Gold (Invitrogen, P36930). Slides were protected from any surrounding light, left to dry overnight and imaged with a Zeiss Axio Scan.Z1.

**AAV expression**. To verify AAV injection sites, animals who had received stereotaxic DREADD, GCaMP8s, or CHR2 injections were sacrificed at ≤18 weeks. Mice were deeply anesthetized with a lethal amount of isoflurane and perfused with 4% PFA. Extracted brains were then stored in 4% PFA overnight at 4°C before being sliced at 60 µm using a vibratome (Leica VT1000S). PVN regions were identified via the Allen Brain Atlas. PVN slices were stained with a DAPI mounting medium (200 µl/slide, Vectashield H-2000) and scanned using a Zeiss Axio Scan.Z1. Images were enhanced and/or pseudo-colored using ImageJ.

### Fiber photometry recordings

**Recording setup**. The Neurophotometrics fiber photometry system (FP3001), operating with Bonsai (Version 2.72) software, was used to acquire photometric data with concurrent video recordings (Arducam, OV9281). Patchcords (MBF Bioscience, NPM-BPC-4) were attached to the fiber ferrules, and held in place by either ceramic sleeves or by a quick-release interconnecter (ThorLabs, ADAL3). Calcium dependent GCaMP fluorescence signals were recorded with light from a 470-nm LED that was band-pass filtered, narrowed with a collimator, reflected by a dichroic mirror, and focused on the sensor of a CCD camera by a 20x objective. To account for auto-fluorescence and possible motion artifacts, a 415-nm LED for calcium-independent isosbestic fluorescence signal was used. The recorded signals were acquired with a sampling rate of 30 Hz for each channel at 100% duty cycle. LED light power was calibrated prior to every recording using a power meter (ThorLabs, PM100USB and S120C). LED light was delivered at a power that resulted in 60µW of 415-nm light, and 85 µW of 470-nm light at the tip of the patchcord.

**Recording conditions.** Photometric recordings occurred in the home cage, with the cage-lid replaced by a custom-fit metal sleeve that extended the cage wall height to 16”. Optic fiber-implanted mice and their pair-housed sister were habituated to the connector and patchcord prior to commencement of experimental schedule. Habituation periods ranged from several days to a week depending on individual animal reaction to the patchcord and to sufficiently aversion train the cagemate. Aversion training was performed during habituation periods by lightly coating the patchcord with diluted capsaicin (EMD Millipore, 211275). The cagemate and implant mouse were observed, and training was repeated until the cagemate associated the unpleasant sensation of capsaicin with chewing of the patchcord. Recordings were approximately 2-hours and were conducted at three different floor temperatures (15C°, 23C° and 29°C), with and without the sister cage-mate. Floor temperature was controlled by placing the home-cage onto a custom aluminum chassis that attached to the Hot/Cold Plate (Ugo Basile, 35300), temperature was then validated by a temperature logging iButton (DS1922L-F5, Embedded Data Systems) placed inside the home-cage. A portion of the nesting material was removed before experiments to prevent occlusion of the animals during the recording (see Supplementary Video 1 for typical amount of bedding left in cage).

**Validation trials using the lactation assay.** To verify the bursting activity of PVN oxytocin neurons observed in photometry recordings, virgin females were crossed with stud males after completing at least two photometry trials (as described above). The dams and pups were housed in their home cages throughout the trial period. Dams were recorded during nursing and lactation. Maternal oxytocin neural activity was recorded at the following timepoints: postpartum day (PPD) 2, 5, 7, 8, 9, and 14. For this experiment the minimum peak amplitude was set to be > 7 standard deviations (sds) above the mean. This threshold was obtained by examining peaks in mothers that clearly displayed pulsatile activity bursts during lactation and closely resembled the peak dynamics that have been previously described during lactation (Yaguchi et al., 2024, 2023; Yukinaga et al., 2022).

**Behavioral characterization.** Videos were uploaded to Noldus Ethovision XT 16, where the physical activity of the mice (calculated total percentage of pixel change between frames) and behaviors were identified. Activity in the paired context was defined as the summed activity for both animals. The behaviors for solo photometry animals were manually scored as follows: Locomotor Activity (LMA), Eating or Drinking (EaDr), Grooming (Groom), Nesting or Nest Building Nest (Nest) where the animal engaged in nest-building behavior or activity within its nest other than grooming, Quiescence (Quies) where the animal is inactive inside its nest, and Stationary (Sta) which was defined as animal inactivity outside of the nest.

The behavior of the photometry animal for paired assays was manually scored as follows: Locomotor Activity (LMA), Eating or Drinking (EaDr), contact initiated (ConI), contact received (ConR), Grooming (Groom), Nesting or Building Nest (Nest) characterized by the photometry animal engaging in nest-building behavior or its activity inside the nest while its cagemate engaged in nest-building behavior, Active huddle (AHud) defined as physical contact and activity between both animals in the nest, Quiescent Huddle (QHud) referred to physical contact and inactivity between both animals in the nest, Quiescence (Quies) where the photometry animal was engaging in inactivity alone in the nest, and Stationary (Sta).

Behavior and activity data was exported at 10 frames per second as a .csv file and imported to R for analysis.

**Calcium data preprocessing**. Fluorescent signal data preprocessing was done with a custom R script as follows: 415 and 470 channels de-interleaved; 470 channel filtered to remove high-frequency noise using a 4th-order Butterworth filter (Van Boxtel et al., 2021) modified to remove end-effect transients; trimmed to remove approximately the first six minutes of data (to mitigate the effect of photobleaching); if necessary, further trimmed to remove effects of disruptions to the patchcord (chewing or twisting); smoothed with a sliding window function (rollapply; width = 9) (Zeileis et al., 2004).

To correct the calcium-dependent signal from effects of photobleaching and heat-mediated LED decay, we first extracted the “fitted values” from the linear relationship between the 470 (calcium-dependent) and 415 (calcium independent) fluorescent signals. Next, to calculate dF/F we divided the 470 signal by these fitted values. Because we observed occasional artifactual shifts in baseline dF/F values, we baseline corrected dF/F using the iterative least squares method (Liland and Mevik, 2011). The Z-scored dF/F was calculated as Z = (χ - μ)/ σ, where μ is the mean dF/F and σ is the standard deviation of dF/F. To define peaks, we used the findpeaks function (Borchers, 2011), with the minimum peak distance set at two seconds and the minimum peak height set at six standard deviations above the mean dF/F. Photometric data were aligned to behavioral data (exported from Noldus Ethovision) and Tb data (exported from Mercury) using common centisecond-resolution timestamps and reduced to a sampling frequency of 10 frames per second.

Peri-event data oriented on specific events (i.e., calcium peaks or start/stop frames of behavioral bouts) were extracted using a custom function that (1) extracts the indices of the events and (2) generates a list of vectors for a specified number of rows before and after the event.

To compare longitudinal physiological patterns of activity in PVN^OT^ neurons in virgin females (primary subjects of this study) compared to nursing and lactation, we made the following modifications to the protocol as described above. First, to quantify PVN^OT^ peak characteristics (amplitude, full width half maximum, interpeak interval) before linear normalization and baseline correction, (i.e., to better describe the dynamics of their native characteristics), we corrected the calcium-dependent signal from effects of photobleaching and heat-mediated LED decay using an exponential decay model. Specifically, we used a self-starting nls (nonlinear least squares) function in R to fit the isosbestic data with the following model: y(t) ∼ SSasymp(timeS, yf, y0, log_alpha), where the measured isosbestic value y starts at y0 and decays towards yf at a rate α. SSasymp is a shortcut that guesses its own parameters; instead of fitting the rate constant α directly, it searches for the logarithm of α: y(t)∼yf+(y0−yf)e^−exp(logα)t. We then linearly scaled this fitted decay to the calcium-dependent data using robust regression (MASS::rlm with the psi function set to bisquare); finally, we divided the calcium-dependent data by this scaled fit to get a corrected signal.

In accordance with previous work, we examined three epochs in the same animals: virgin females; early-stage lactation (PPD 2-7); and late-stage lactation (PPD 8-14). To calculate the full-width half maximum (FWHM) of PVN^OT^ peaks, we used a custom function that performs the following steps: finds each peak; determines the half-max value; interpolates where the signal crosses the half-max value; computes the FWHM as the difference between these crossing points. To enable visual comparison of waveform width across groups in dependent of amplitude differences, we derived peak-normalized average waveforms using a normalization procedure for every peak prior to averaging.

Specifically, for each peak we (1) baseline-subtracted the trace by subtracting the mean fluorescence in a pre-peak baseline window, and then (2) divided the baseline-subtracted waveform by its own maximum value to scale the event amplitude to 1. We then computed the mean ± SEM of these peak-normalized waveforms across events within each group. To calculate the interpeak interval (IPI), we used the lead function in R to compute the time (in seconds) from one peak to the next on a per-recording basis.

### Computer vision analysis

**Processing Flir .seq files.** RGB videos, timestamps, and thermal data were extracted from Flir thermographic .seq files using a Python-based adaptation of *ThermImageJ* ^3^. Raw images were obtained via Flirpy ^4^, converted to RGB, and merged into AVI video files. Metadata, including timestamps and frame rates, was extracted using *Exiftool* ^5^. This workflow provided thermal images for model training, AVI videos for analysis, and timestamps and temperature data for integration with behavioral information. The processing script is available at https://github.com/j-landen/seq_process.

**Skeleton-Guided Bodypart Segmentation (SGBS) for thermographic temperature extraction.** To quantify region-specific surface temperature from FLIR thermography videos in awake mice, we developed Skeleton-Guided Bodypart Segmentation (SGBS), a two-stage computer vision pipeline that combines DeepLabCut (DLC) keypoint tracking with an instance-segmentation network (Mask R-CNN) (code: https://github.com/j-landen/SGBS). First, a DLC model was trained on 1,800 manually labeled frames to detect 10 dorsal keypoints (nose, head tip, implant, neck base, center back, tail base, mid-tail, patchcord, and left/right shoulder). For each thermography frame, these keypoints were converted to a set of per-keypoint spatial heatmaps and fused with the thermal image as additional channels that guide segmentation (i.e., a skeleton-conditioned attention/fusion step that biases the network toward anatomically plausible regions even when posture, bedding, or patchcords alter thermal texture). Second, a custom Mask R-CNN model was trained on 457 manually annotated images (Darwin/V7) to segment three regions of interest: interscapular BAT, rump, and the full dorsal surface.

Model performance was evaluated on a held-out validation set not used for training (20% of annotated images), using standard segmentation accuracy metrics (per-class intersection-over-union and Dice/F1) and by visual inspection of predicted masks overlaid on raw frames. As an additional control, we trained an unmodified Mask R-CNN (same backbone initialization, same training/validation split, and identical training schedule) and compared loss curves and validation performance between the baseline and skeleton-guided models. Loss is calculated as the sum of three components: (1) a classification loss (cross-entropy) that assesses how accurately each body part is identified, (2) a bounding box regression loss that measures how close the predicted box is to the ground truth, and (3) a mask loss (binary cross-entropy) that evaluates pixel-by-pixel agreement between the predicted mask and the ground truth mask. Finally, masks were mapped to radiometric pixel values to extract per-frame temperature statistics (mean, min, max, SD) for each region. To establish physiological (external) validity independent of segmentation training, SGBS-derived temperature trajectories were temporally aligned with simultaneously recorded core body temperature from implanted thermologgers; thermologger data were not used for model training. SGBS measures showed strong positive correspondence with core temperature dynamics and reproduced expected temporal relationships among BAT, dorsal, and rump signals (e.g., BAT changes preceding core temperature, consistent with thermogenic physiology).

### Optogenetic stimulation

**Recording setup**. Blue light (450 nm) stimulation was delivered using the Neurophotometrics fiber photometry system (FP3001), with simultaneous video and thermal recording. Mono-filament patchcords (Doric Lenses, MFP_200/220/900-0.37_3m_FC-MF1.25) were attached to the fiber ferrules and held in place by a quick-release interconnecter (ThorLabs, ADAL3). Optogenetic laser power was measured prior to every recording using a power meter (ThorLabs, PM100USB and S120C). Optogenetic activation used a frequency of 10 Hz, with a 20 ms pulse width (20% duty cycle) and 10 mW of power measured at the tip for either 10- or 20-second trains.

Optogenetic recordings occurred in the home cage, with the same setup as fiber photometry recordings, described above. Mice were pair-housed until recording. One animal was tested at a time, while the other animal was placed in a holding cage for the duration of the experiment. During experiments, animals were observed remotely from a different room. The human observer turned on blue light stimulation after the animal had been inactive for a minimum of two minutes. Stimulations were a minimum of ten minutes apart from one another. Experiments were two hours long, averaging four to seven stimulations. Every animal was tested at least twice using each stimulation time (10s and 20s), until a minimum of 10 total stimulations occurred per animal, per stimulation time.

**Optogenetic data preprocessing.** For each stimulation, data spanning 5 min (–300 s) before to 10 min (+600 s) after light onset were extracted, with the onset designated as time 0. Core body temperature (Tb) data was recorded once every five seconds for the duration of the experiment. We therefore subset all other data to the same rate for core Tb analyses.

Activity levels were measured at a rate of 10 Hz, so all data were subset to the same rate for activity analyses. As activity is measured using percent pixel movement over the entire image, activity was first set to a logarithmic scale, to measure small differences in change, then normalized using a Z score across mouse IDs to account for differences in baseline activity levels.

Feature-specific surface temperatures using SGBS were measured at a rate of around 20-30 Hz, depending on the Flir A50’s output. As previously described, all surface temperature data was subset to 10 Hz and aligned to surface temperature data based on the nearest timestamps when the laser was turned on (< 0.1s deviation). To eliminate the effect of outliers originating from an incorrect segmentation output, all surface temperature data were smoothed with a rolling average over one half of a second. To eliminate differences in baselines per animal, every value was baseline-normalized using ΔT/T, where each temperature value was expressed as a change from the pre-stimulation mean (3-minutes prior to light on) relative to that baseline.

To assess whether optogenetic effects varied with initial thermal state, core body temperature at the onset of each blue-light pulse (Tb-T0) was recorded. Because Tb-T0 values were generally higher than those observed during spontaneous PVNOT calcium peaks, stimulations were stratified for each mouse into “low Tb-T0” and “high Tb-T0” trials using that animal’s mean Tb-T0 as the cutoff. The two strata were then analyzed separately.

### Statistics

**Statistical results are reported in Table S1.**

**Effect of behavior state on FOS expression.** To determine the effect of behavior on FOS expression, brain ROIs (divided into left and right sides, where appropriate) were scanned by CellProfiler 4.2.1 to determine the total number of cells, as marked by DAPI. DAPI ROIs were shrunk to a point and then expanded by a factor of 6, and classified as being either FOS-positive or FOS-negative, as marked by presence of the secondary antibody (Alexa Fluor 555), using the same parameters for every slice. These per-ROI portions were then standardized using a Z-score between different histology days, to account for any batch effects of antibody binding or fluorescence. A linear regression model was used on a per-ROI basis, and a post-hoc tukey test was used to determine the effect of behavior on FOS expression, for each brain region. P-values were adjusted for multiple comparisions using the Holm method.

**Relationship between PVN^OT^ calcium activity and behavior state, activity, and body temperature**. To determine the effect of different behavioral states on calcium peak amplitude and frequency, we fit LMMs with amplitude or frequency as the dependent variable, behavioral state as the independent variable, and mouseID as a random effect. These models were run for both solo and paired conditions.

To determine the effect of social context and floor temperature on calcium peak amplitude and frequency, we fit LMMs with amplitude or frequency as the dependent variable, social context or floor temperature as the independent variable, and mouseID as a random effect.

To determine how calcium peak frequency is affected by behavioral state bout length (i.e., for quiescence and quiescent-huddle) and floor temperature, we fit a LMM with per-bout peak counts as the dependent variable; bout-length, floor temperature, and their interactions as the independent variables; and mouseID as the random effect. We subsequently extracted the standardized beta coefficients using the sjPlot package (Lüdecke, 2013).

To determine how locomotor activity changed during a 300 sec. interval of time surrounding calcium peaks (i.e., peri-peak time), we first z-scored the activity data (estimated at 10 Hz), then smoothed it according to the mean using the rollapply function (Zeileis et al., 2004), and finally calculated the per-second mean for each individual. Next, we calculated normalized activity data across the 600 sec interval, including the mean normalized activity before and after each peak (i.e., “before-after”). Last, we fit a LMM with normalized activity as the dependent variable, “before-after” as the independent variable, and mouse-ID as a random effect.

To determine peak probability during defined behaviors we used logistic regression. First, we used custom R code to (1) extract the onset and offset of behavioral bouts, and (2) label rows of data (at 10 frames per second) according to whether they occurred within a minute containing (or not containing) a calcium peak (i.e., PeakMinute). Next, we fit a binomial generalized linear model with PeakMinute as the dependent variable; the independent variable was the time span before, during, and after the behavioral epoch of interest (e.g., quiescence offset). Next, we extracted the per-frame predicted values (including standard error) from this model.

To compare the statistical distributions of Tb during calcium peaks *vs*. baseline, we plotted Tb according to the PeakMinute designation described above. We then used a LMM with Tb as the dependent variable; the independent variables were PeakMinute and mouseID as a random effect. To determine whether PVN^OT^ peaks were associated with enhanced thermogenic changes at rest-to-active transitions, we performed transition-aligned analyses of body temperature followed by linear mixed modeling. First, we extracted 600 sec peri-transition segments (−300 to +300 sec; sampled at 10 Hz) aligned to the offset of quiescence and quiescent-huddle bouts. Each transition was categorized as Peak+ if one or more PVN^OT^ peaks occurred within the 100 sec interval preceding bout offset (−100 to 0 sec), and Peak− otherwise. To account for inter-individual variability in Tb set point, temperature was z-scored within mouse prior to analysis. For each transition, we quantified the post-offset change in scaled body temperature (Δz-Tb) by subtracting a pre-offset baseline (mean z-Tb from −60 to 0 sec) from the mean post-offset value (mean z-Tb from 0 to 300 sec). We then fit separate LMMs for quiescence offsets and quiescent-huddle offsets with Δz-Tb as the dependent variable, PeakPlus (Peak+ vs. Peak−) as the independent variable, and mouseID as a random effect.

**PVN^OT^ peak characteristics: before mating *vs*. early/late stages of lactation**. To determine the effect of life-history phase (virgin, early-stage and late-stage lactation) on peak amplitude, FWHM, and interpeak interval we used LMMs with epoch (PPD 2-7; PPD 8-14; virgin) as a fixed effect and mouseID as a random effect to control for repeated measures. We performed posthoc tukey comparisons for each pairwise combination of fixed effect level using the emmeans package.

**Effect of optogenetic stimulation on thermal profile.** To determine the effect of light stimulation on Tb in ChR2+ and light-only controls separately, Tb data were binned per minute, with each value representing the mean of the 60 seconds following. We used a linear mixed-effects model (LMM) with Tb as the dependent variable, the interaction between light-status and minute as independent variables, and mouseID as a random effect. A post-hoc tukey test was used to determine per-minute differences.

To determine the effect of light stimulation on physical activity in ChR2+ vs. light-only controls separately, activity was binned per minute, and we used the same LMM as above, but with activity as the dependent variable.

To determine the effect of light stimulation on surface temperatures (BAT, rump, or dorsal surface) in ChR2+ vs. light-only controls separately, each surface temperature was analyzed separately, first by binning temps per minute, and then using the same LMM as above, with surface temps as the dependent variable.

To further determine the effect of light stimulation on direction of temperature change, slopes before and after light onset were analyzed with a second LMM that included fixed effects of time, proximity to stimulation (pre vs. post), and their interaction, plus a random intercept for mouse ID; frame-level weights were set to the inverse of each frame’s temperature SD. A significant interaction term indicated that temperature trajectories differed between ChR2+ and control groups across the stimulation boundary.

To assess the effect of light stimulation on behavior in ChR2+ versus light-only controls, we quantified cumulative time spent in each behavioral state following stimulation. First, linear regression models were applied to individual arousal behaviors to evaluate their contribution to group differences. As no single behavior showed a significant effect, all arousal behaviors were grouped, and a second linear regression model was used to assess whether total arousal time differed between groups.

To quantify the temporal progression by which optogenetic stimulation influenced core body temperature while accounting for concomitant changes in locomotor activity and BAT surface temperature, we used lagged regression and cross-correlation analyses of thermal dynamics. First, we performed lagged regression analyses on the derivative of core Tb (dTb/dt). For these analyses, frame-level data were first restricted to a peri-stimulus window (time-from-stimulation onset: −270 to +570 s) and then aggregated into 1 s bins within each trial. Within each trial, we computed first differences for Tb and BAT surface temperature (dTb(t) = Tb(t) − Tb(t−1); dBAT(t) = BAT(t) − BAT(t−1)). Photostimulation was represented as a binary 1 s time series (stim(t) = 1 during seconds containing any LightOn frames, else 0) and transformed into an impulse-response regressor using a kernel (τ = 30 s), producing a decaying stimulation effect term (stim_k(τ)). We then fit a LMM, “mod_nonstim”, with dTb as the dependent variable and fixed effects including stim_k(t), distributed lags of activity (Activity(t−L)) and BAT derivative (dBAT(t−L)) over L = 0–60 s (10 s increments), and random intercepts for mouse ID and trial. Predictors were z-scored to facilitate model convergence and comparison of effect sizes. (mod_nostim: dTb_z ∼ Act_L0 + Act_L10 + Act_L20 + Act_L30 + Act_L40 + Act_L50 + Act_L60 + dBAT_L0 + dBAT_L10 + dBAT_L20 + dBAT_L30 + dBAT_L40 + dBAT_L50 + dBAT_L60 + (1 | mouseID) + (1 | trial)).

To test whether stimulation explained variance in dTb beyond that captured by activity and BAT surface dynamics, we compared the full mod_nonstim model to an otherwise identical model excluding the stimulation parameter (stim_k(τ)) using a likelihood-ratio test.

To assess temporal ordering of changes in BAT surface temperature and activity relative to changes in core temperature, we computed trial-wise cross-correlation functions between derivative time series (dBAT vs. dTb; Activity derivative vs. dTb) within the peri-stimulus window, using a lag search range of ±180 s. For each trial, we extracted the lag corresponding to the peak cross-correlation coefficient (peak-lag), where positive lags indicate that the predictor time series precedes dTb. Peak-lags were summarized across trials within each mouse.

---

**Table S1.**
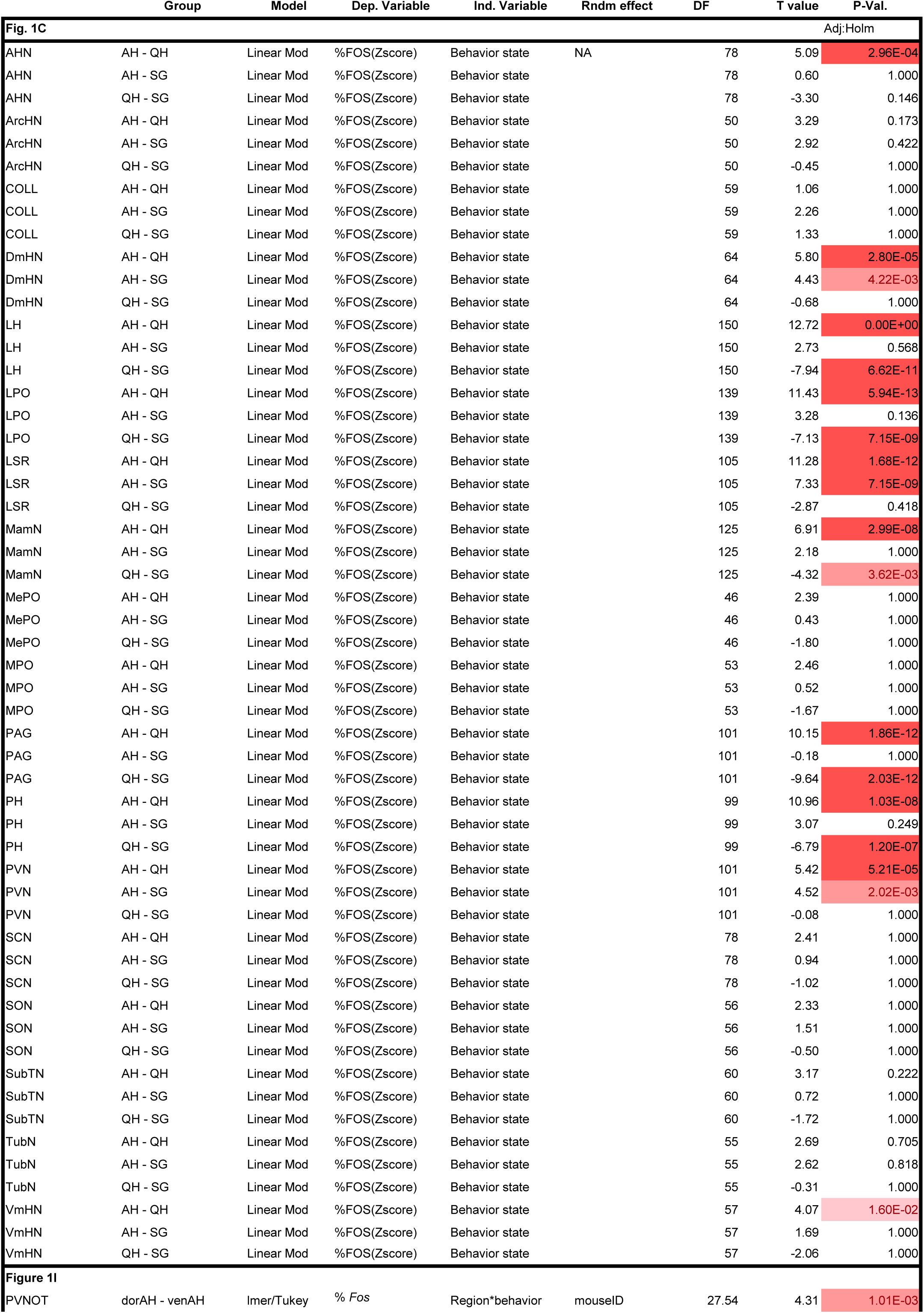
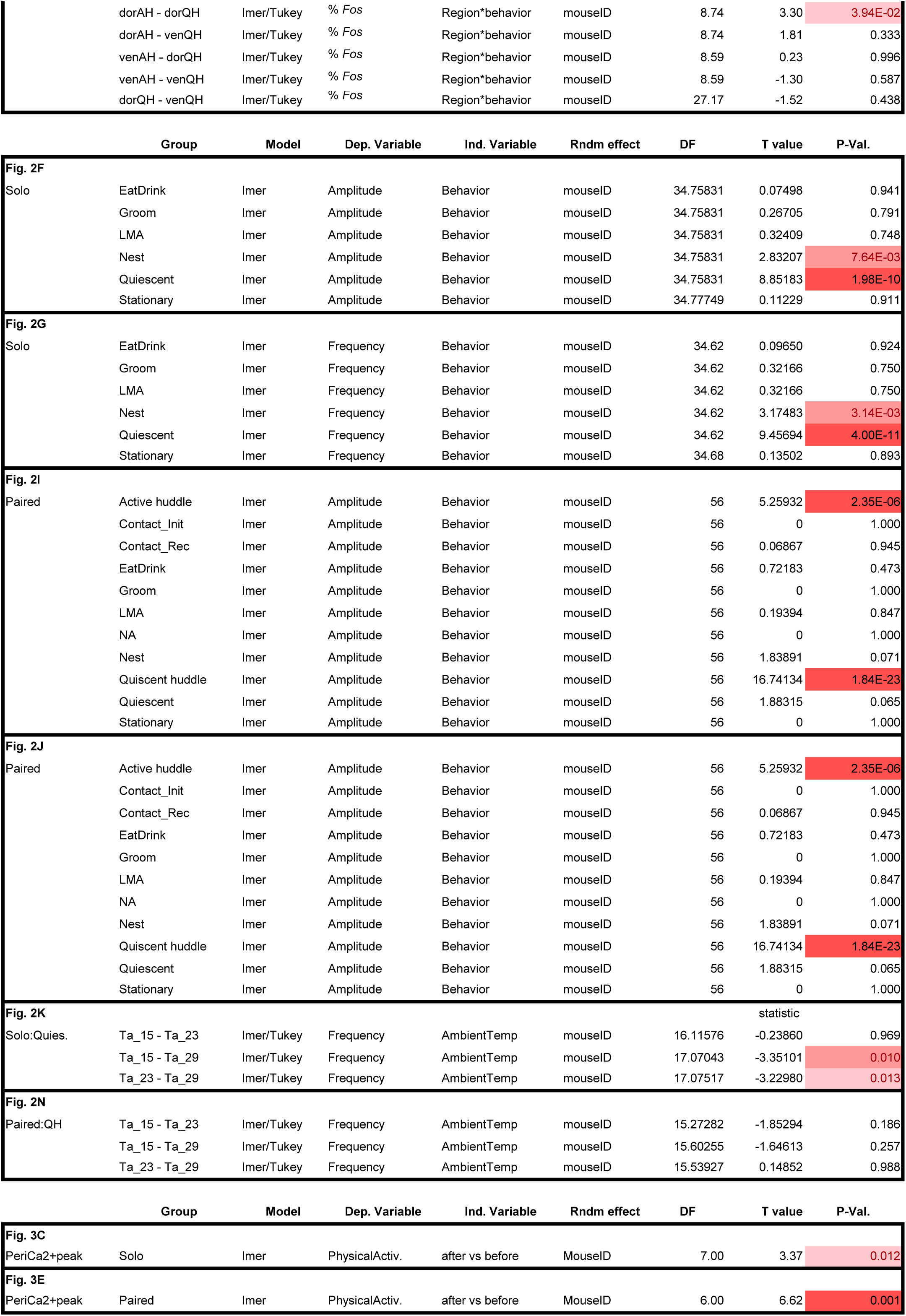
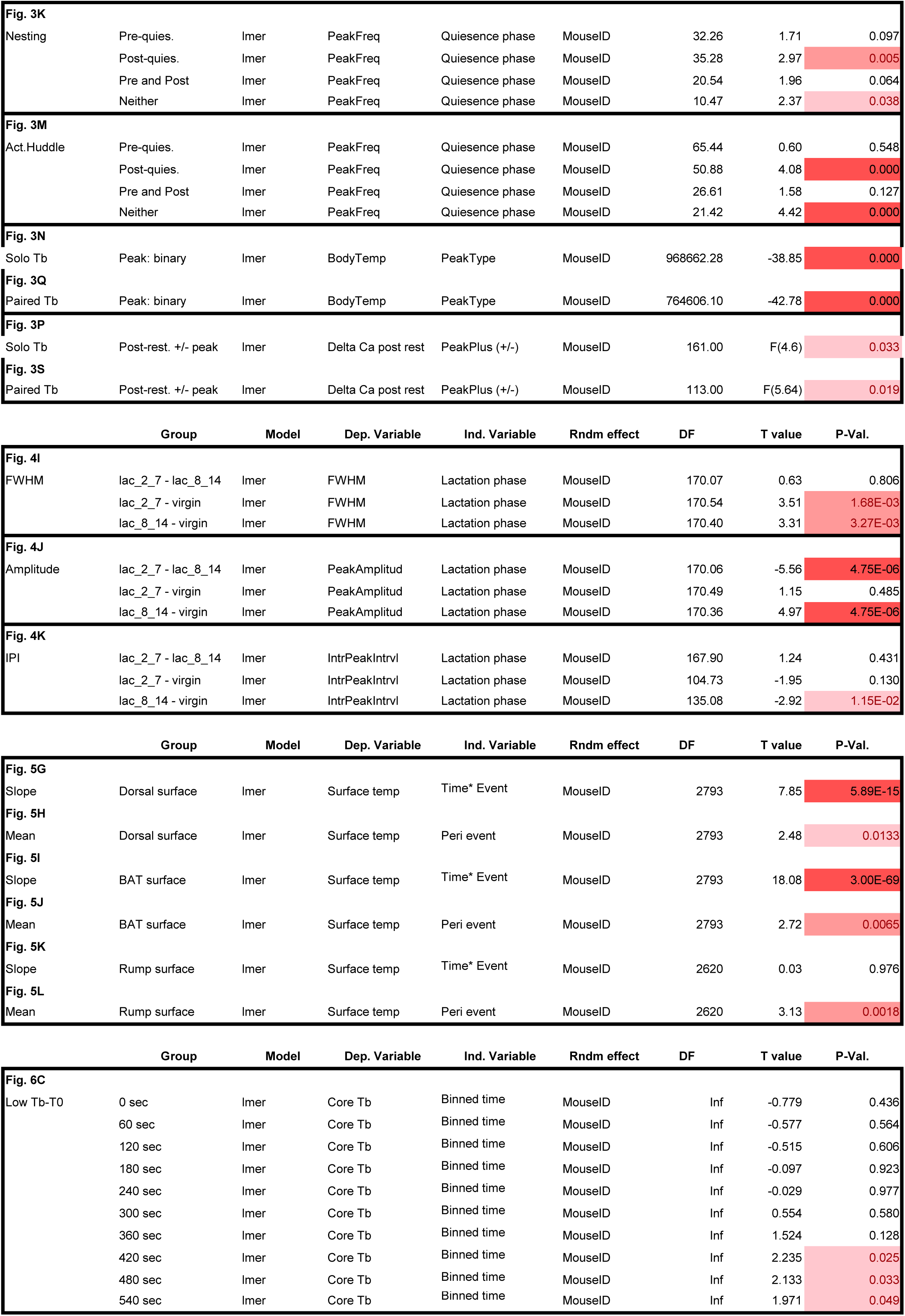
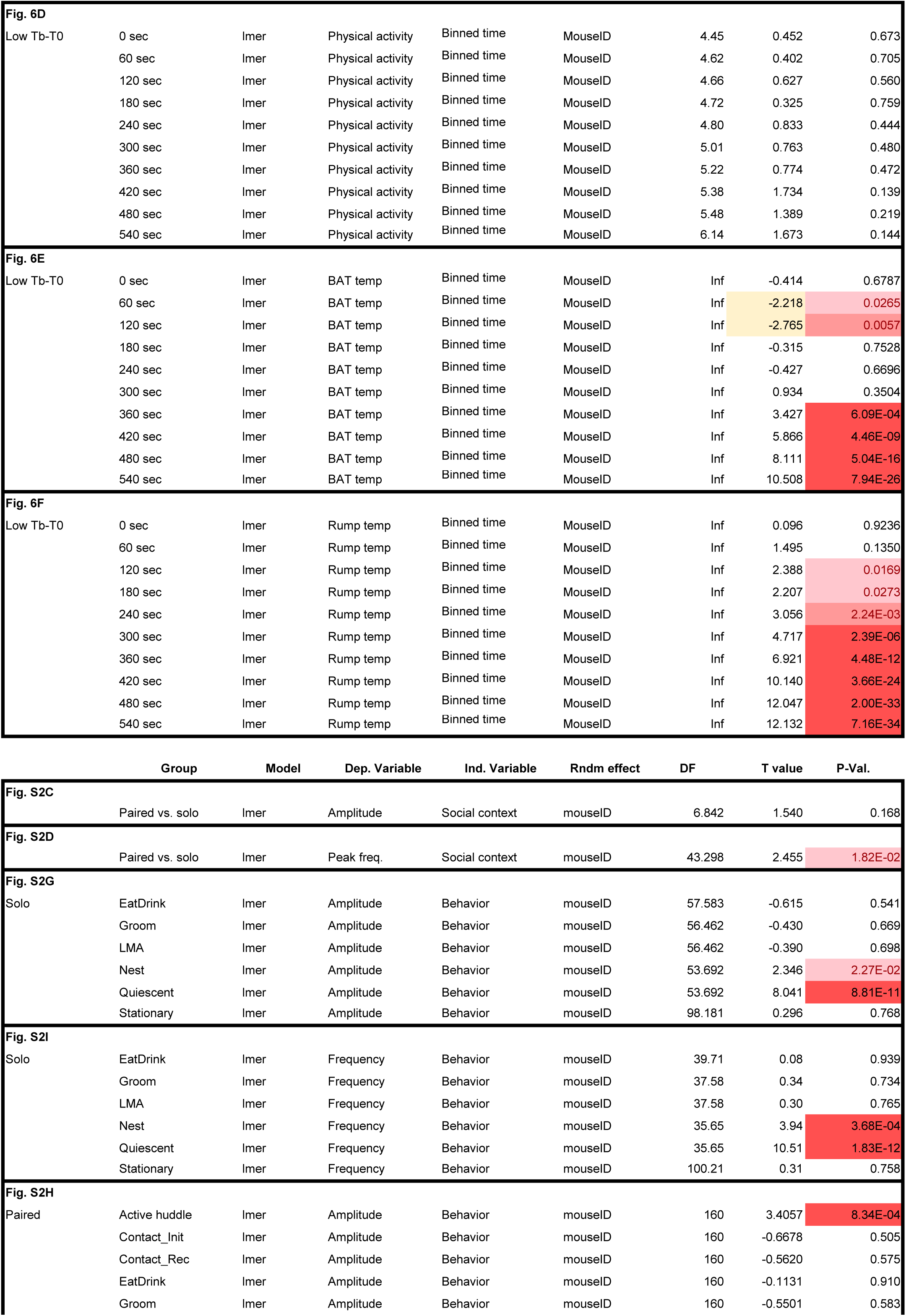
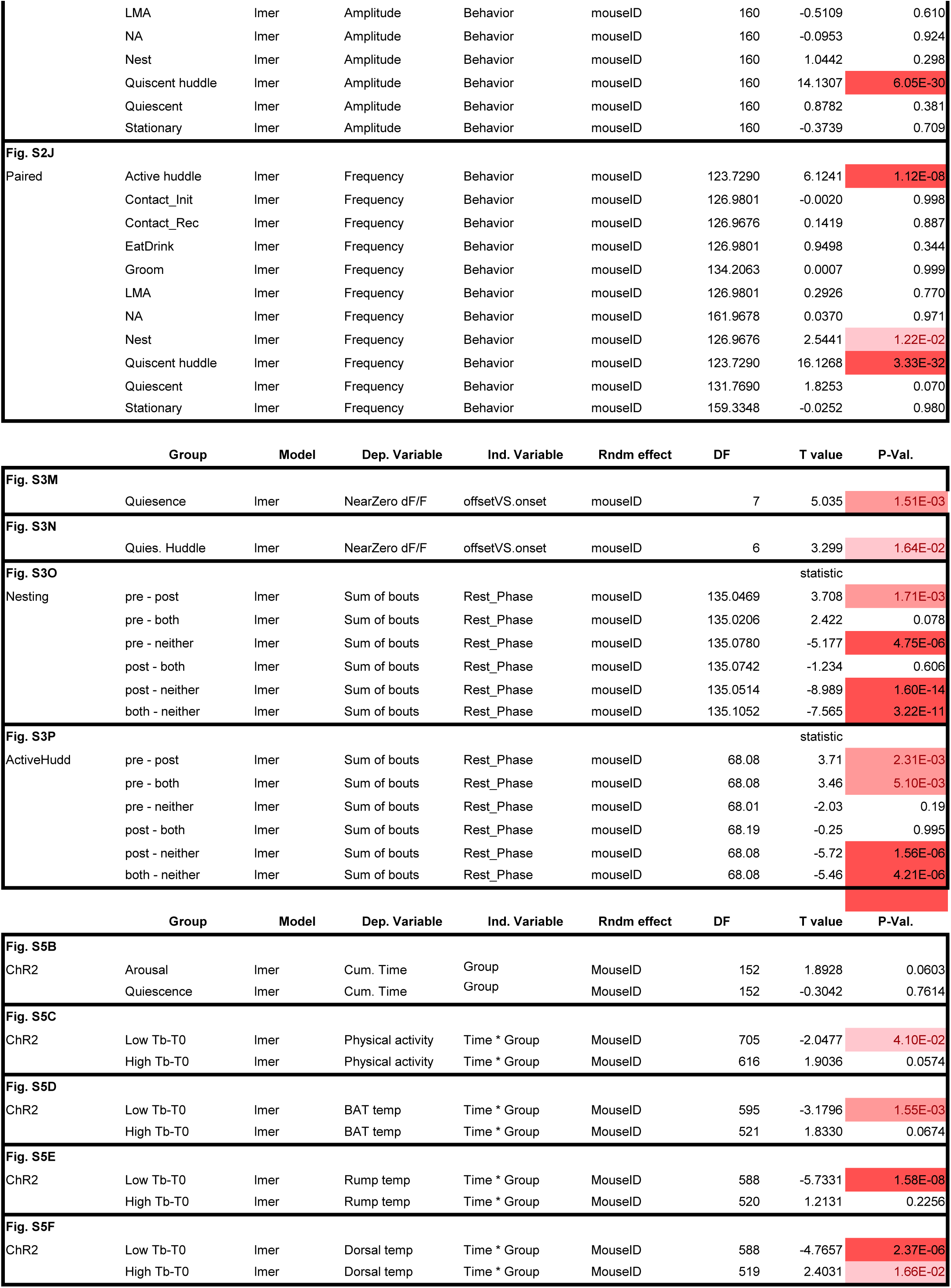

## Supplementary Figures

**Figure S1.**
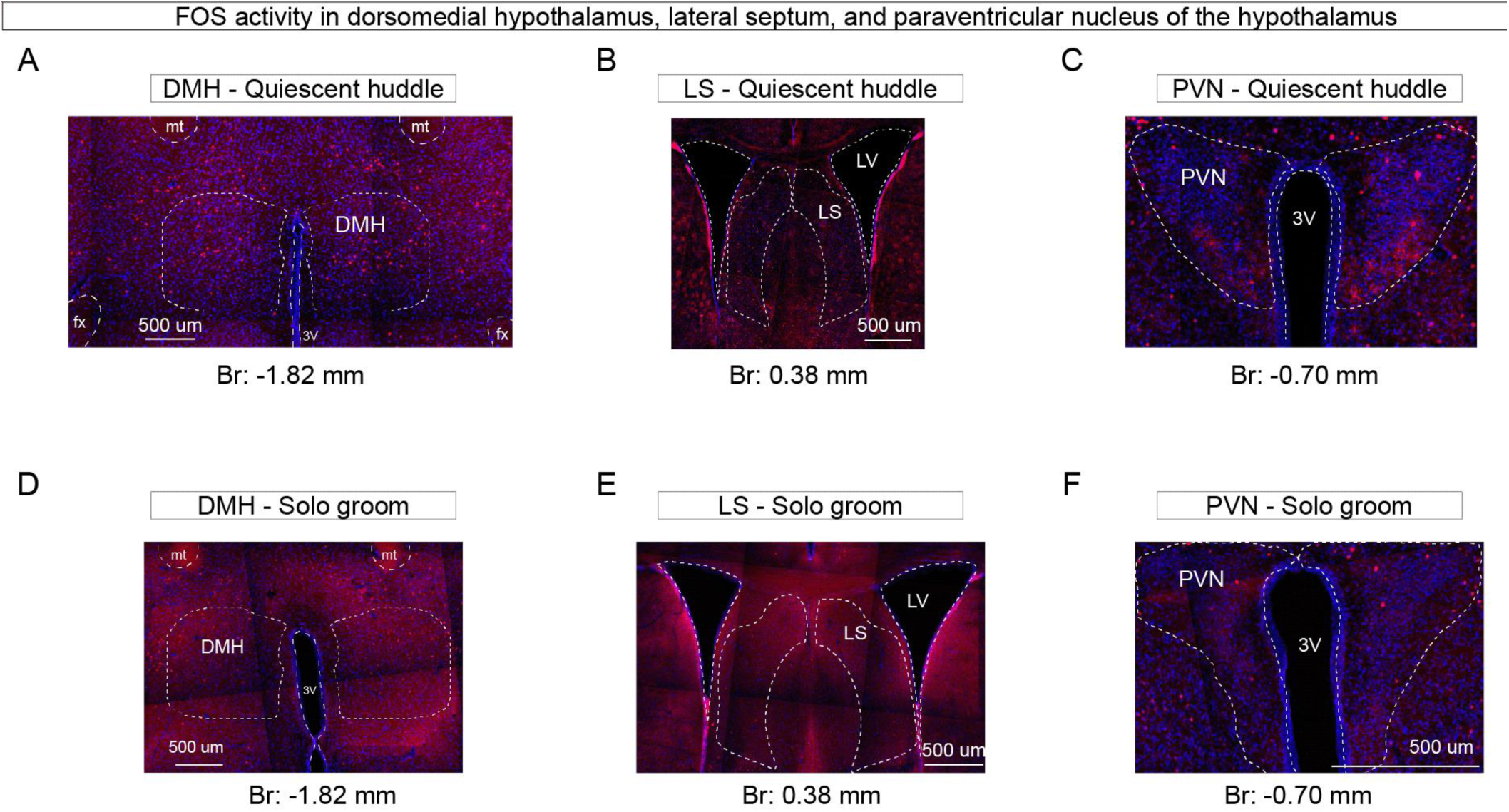
Histology of FOS activity in the DMH, LS, and PVN. **Related to Figure 1.** **(A - C)** Representative histology images showing quiescent huddling associated FOS expression in the DMH **(A)**, LS **(B)**, and PVN **(C)**. **(D – E)** Representative histology showing solo grooming associated FOS expression in the DMH **(D)**, LS **(E)**, and PVN **(F)**. Abbreviations: 3V: third ventricle; DMH dorsomedial hypothalamus; LS lateral septum; mt: mammillothalamic tract; PVN paraventricular nucleus of the hypothalamus.

**Figure S2.**
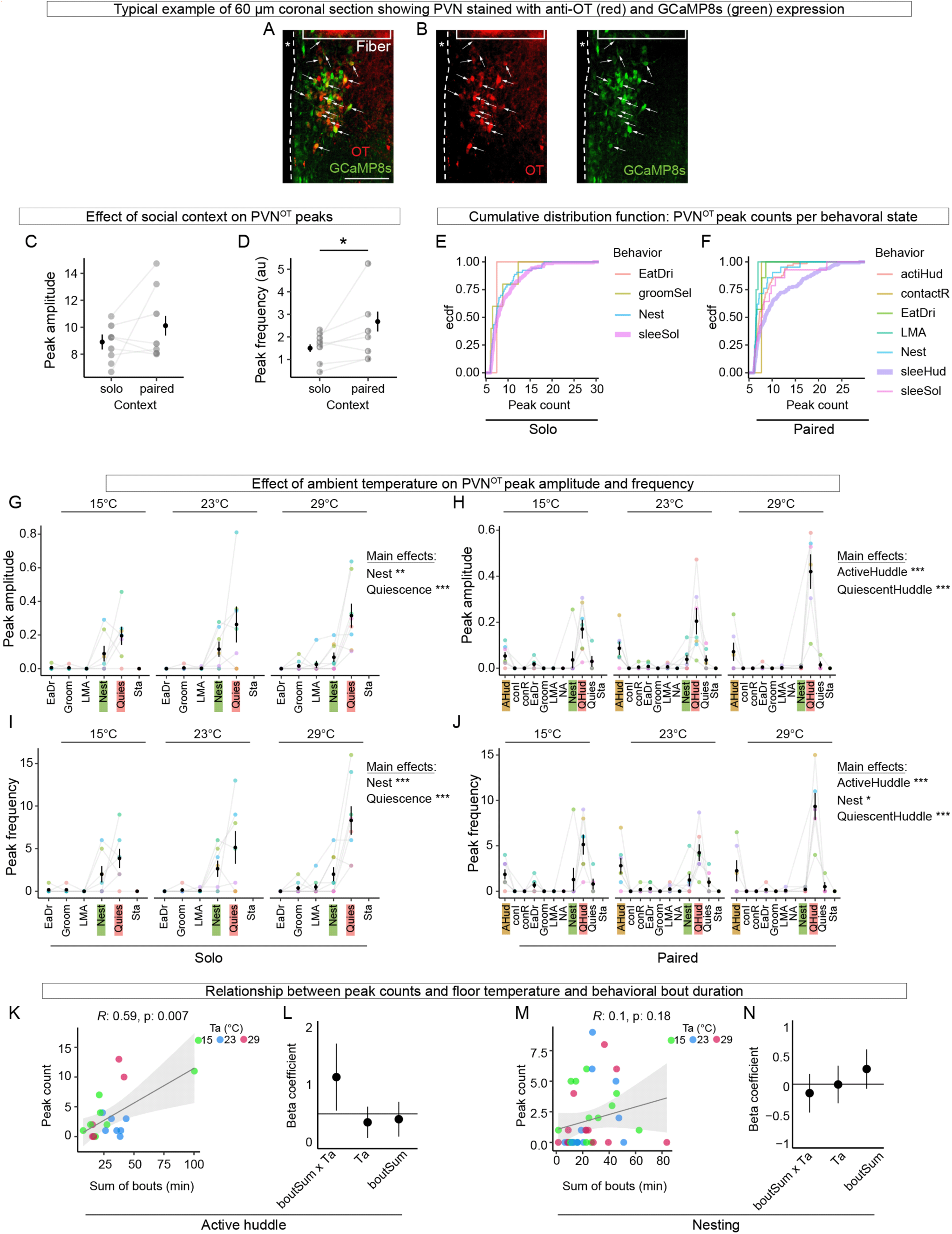
Characterization of PVN^OT^ Ca^2+^ peaks and their associations with behavioral states. **Related to Figure 2.** **(A - B)** Representative coronal brain slice showing the location of the optical fiber and expression of oxytocin peptide (red) and GCaMP8s (green) in the PVN. Arrows indicate some cells co-labeled with GCaMP8s and anti-OT. The third ventricle is designated by “*”. Scale bar 100 μm. **(C - D)** Effect of social context on PVN^OT^ peak amplitude **(C)** and frequency **(D)**. **(E - F)** PVN^OT^ empirical cumulative distribution functions (ecdf) of peak counts according to behavioral state in solo **(E)** and paired **(F)** conditions. **(G - H)** Effect of floor temperature on PVN^OT^ peak amplitude. Relationship between floor temperature and behavior state in solo **(F)** and paired **(G)** females. **(I - J)** Effect of floor temperature on PVN^OT^ peak frequency. Relationship between floor temperature and behavior state in solo **(I)** and paired **(J)** females. **(K - N)** Effect of floor temperature and active behaviors on PVN^OT^ peak counts for active huddling **(K-L)** and nesting **(M-N)**. Peak count according to total duration of bouts per trial; *R* and p values shown at top **(K,M)**. Beta coefficients from a model of the effect of floor temperature, total duration of bouts (boutSum), and the Ta*boutSum interaction on peak count **(L,N)**. Statistical results are from linear mixed models. N = 8 solo, N = 7 paired, N = 50 recordings. Data shows mean ±SEM. P < 0.05 *, P < 0.01 **, P < 0.001 ***.

**Figure S3.**
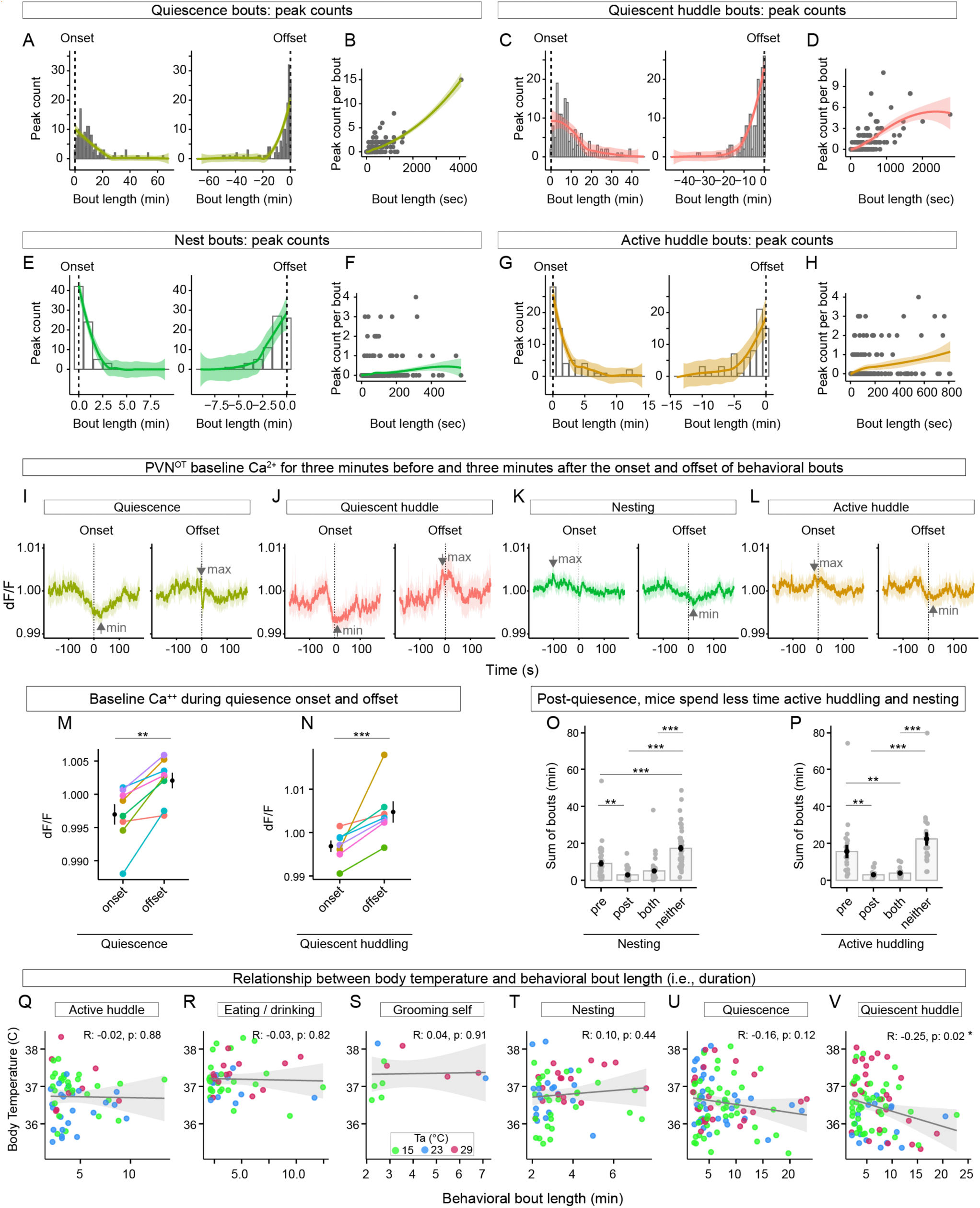
PVN^OT^ Ca^2+^ peaks during behavioral states and transitions. **Related to Figure 3**. **(A - D)** Peak counts for bouts of resting behaviors. Quiescence onset/offset peak counts **(A)** and peak count per bout **(B)**. Quiescence huddle onset/offset peaks counts **(C)** and relationship between bout length and peak count **(D)**. **(E - H)** Peak counts for bouts of active behaviors. Nesting onset/offset peaks counts **(E)** and relationship between bout length and peak count **(F)**. Active huddle onset/offset peak counts **(G)** and relationship between bout length and peak count **(H)**. **(I-L)** Extended analysis of baseline PVN^OT^ Ca^2+^ surrounding bout onsets and offsets. For each behavior, the minimum and maximum dF/F is indicated with an arrow. **(M - N)** Per-individual means of Ca^++^ dF/F during onset and offset (i.e., near-zero values) of two resting behaviors: quiescence **(M)** and quiescent huddle **(N)**. **(O - P)** Sum of bouts according to the phase of quiescence. “Neither” refers to bouts that did not adjoin bouts of quiescence. Nesting bouts **(O)**. Active huddling bouts **(P)**. Post quiescence nesting and active huddling is relatively rare. **(Q-V)** Regression analysis of bout length and Tb. Bouts <2min are excluded. Shown are the Pearson correlation (*R*) and associated p-value from a linear regression. Length of quiescent huddle bouts is negatively correlated with body temperature **(N)**. **M-P**: linear mixed models; N = 8 mice/50 recordings. **(Q-V): linear model;** N = 6 mice/27 recordings. All data shows mean ±SEM. P < 0.05 *, P < 0.01 **, P < 0.001 ***. Full statistical analysis in Table S1.

**Figure S4.**
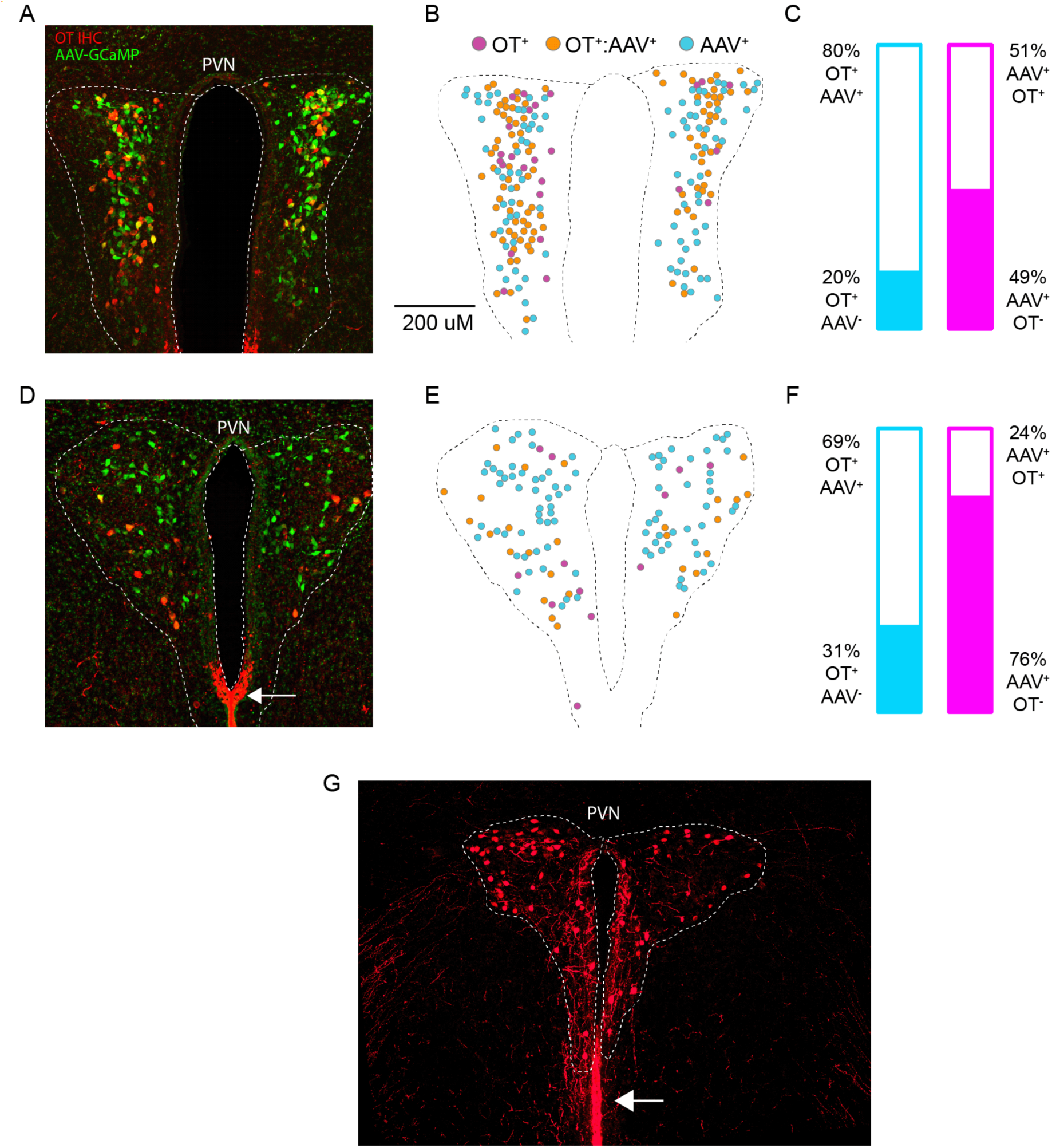
Intra-individual distribution and quantification of OT immunoreactivity and Cre-dependent GCaMP expression. Related to **Figure 4**. **(A)** The paraventricular hypothalamus (PVN) showing oxytocin immunoreactivity (red) and Cre-dependent expression (green). **(B)** Re-construction of (A) showing the distribution of neurons with OT immunoreactivity, GCaMP expression, and their overlap. **(C)** Quantification of cellular overlap in (B). **(D)** An adjacent slice from the same animal as (A). Strong OT immunoreactivity or processes along the ventral third ventricle (white arrow). **(E)** Re-construction of (D) showing the distribution of neurons with OT immunoreactivity, GCaMP expression, and their overlap. **(F)** Quantification of cellular overlap in (E). **(G)** Image showing OT immunoreactivity in processes near the third ventricle, below the PVN.

**Figure S5.**
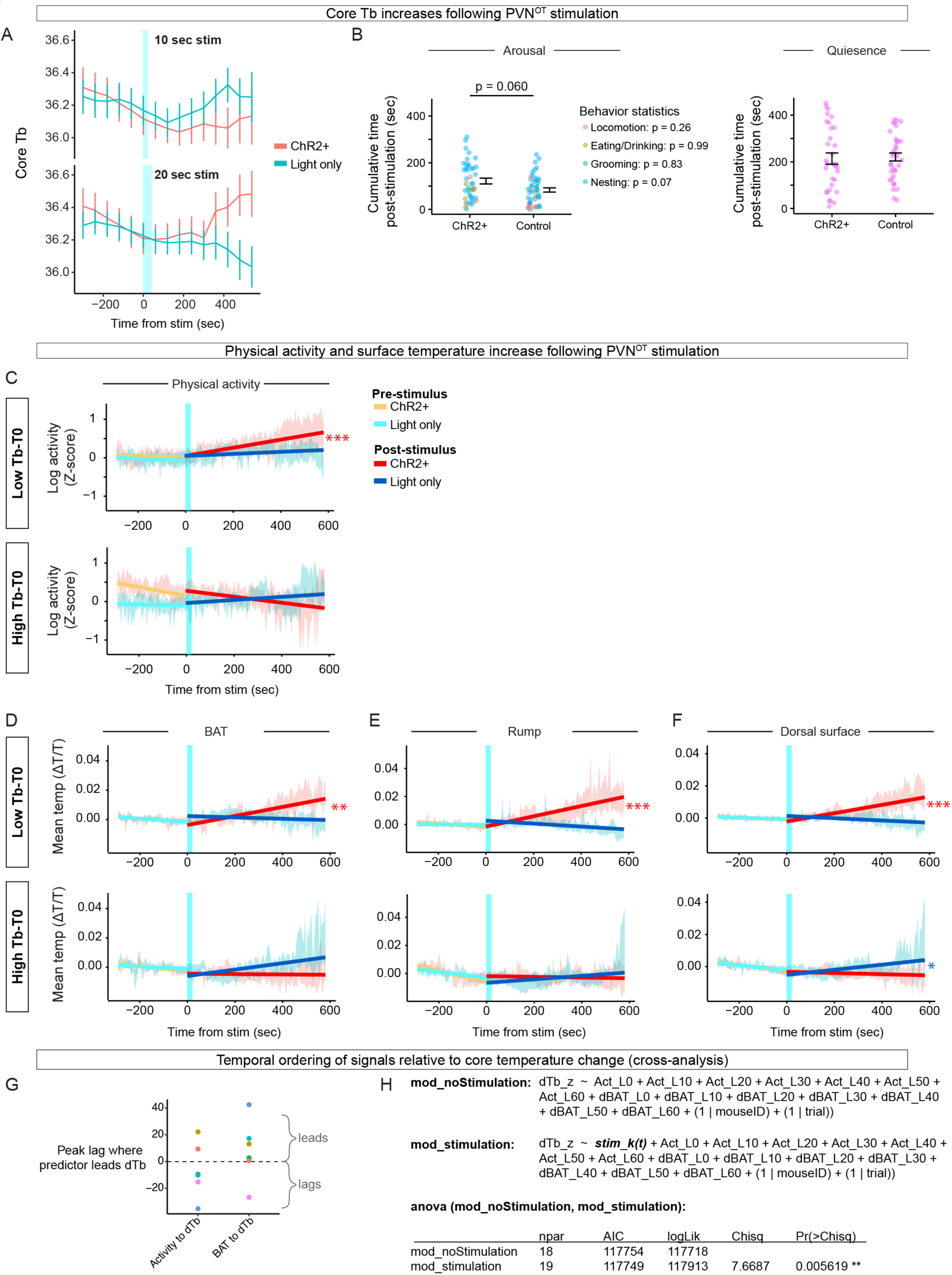
**(A)** Blue light stimulation (time 0; shaded blue rectangle) during 20s vs. 10s stimulations in ChR2+ animals compared to light-only animals. **(B)** Cumulative time spent in arousal/awake (left) and rest/quiescent (right) behaviors following 20s PVNOT stimulation. Each point represents a trial, color-coded by behavior. Black bars show mean ± SEM. **(C)** Linear regression line ± SEM of physical activity (log-transformed Z-score) 5 min before and 10 min following 20s PVNOT stimulation. **(D - F)** Same analysis as in **(C)** for surface temperatures (ΔT/T) of BAT **(D)**, rump **(E)**, and dorsal surface **(F)**. Stimulation in ChR2+ animals significantly increased temperatures in the low Tb-T0 group, with opposite or no effect in the high Tb-T₀ group. P < 0.05 *, P < 0.01 **, P < 0.001 ***. Full statistical analysis in Table S1. **(G)** Mouse-level mean peak cross-correlation lags between dTb/dt and either dBAT/dt (“dBAT to dTb”) or activity (“Activity to dTb”) computed over a ±180 s window. Positive values indicate the predictor precedes dTb/dt; the dashed line marks zero lag. d/dt represents the derivative with respect to time (t). **(H)** Lagged regression model testing the effect of light stimulation on derivative Tb (dTb) while controlling for the effects of activity (Act) and derivative BAT-surface temperature (dBAT). Act and dBAT are represented as lags over 0-60 sec (10 sec increments). In the stimulation model, the light stimulation term (stim_k(t)) is represented as a decaying impulse-response using a 30 sec kernel.

**Figure S6.**
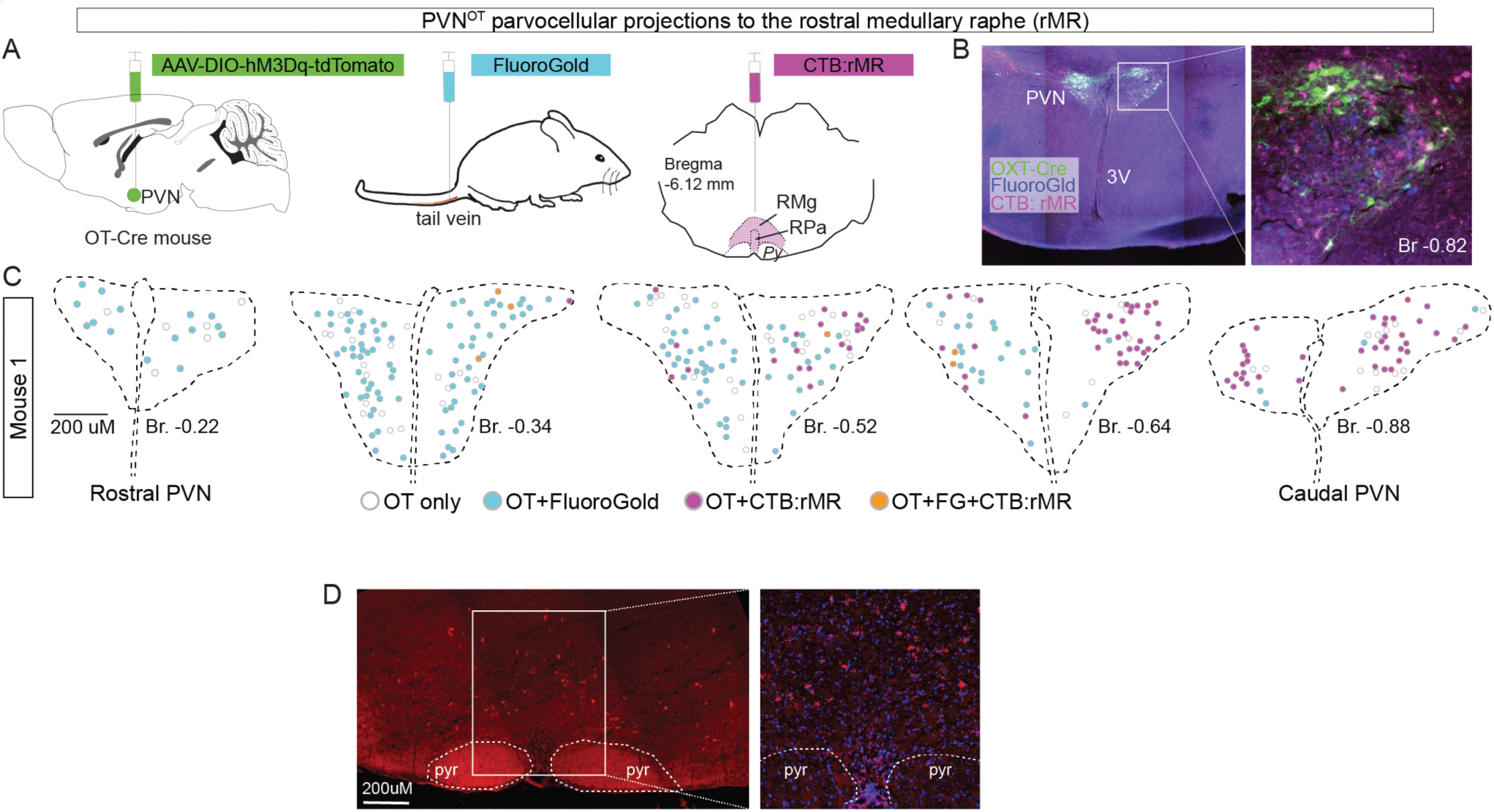
PVN^OT^ parvocellular projections to the rostral medullary raphe. **(A-D)** PVN^OT^ parvocellular projections to the rostral medullary raphe (rMR). Scheme of injections to label magno- and parvo-cellular OT neurons **(A)**. Representative histology showing cells labeled for OXT-Cre, FluoroGold, and CTB **(B)**. Distribution of PVN^OT^ neurons retrogradely labeled with FluoroGold and CTB **(C)**. Each map was made from one coronal section. Cells double-labeled with FluoroGold and OT-Cre were mostly distributed in the rostral part of the PVN; cells double-labeled with CTB and OT-Cre were in the caudal part of the PVN **(C)**. Fluorescent in situ hybridization of OXTR (red) in the rMR region **(D)**.

## Video Legends

**Video S1. Photometry recording of virgin mouse in the homecage**. A large-amplitude PVN^OT^ peak occurs at 0:00:03s. Video is sped up to 4x.

**VideoS2. Photometry recording of a lactating dam in the homecage with pups on PPD 14**. The first large-amplitude PVN^OT^ peaks occur at approximately 0:00:04s. An even larger peak typical of late-stage lactation occurs at 0:00:15s. Video is sped up to 4x.

**VideoS3. SGBS enables real-time segmentation of thermally defined anatomical features in freely moving mice.** Example video shows the output of the SGBS model, which accurately identifies and tracks three anatomical regions – BAT, rump, and dorsal surface – in a freely moving mouse in the homecage using thermal imaging. Video is sped up to 4x.

